# Cohesin promotes genomic stability by suppressing unequal sister chromatid exchange

**DOI:** 10.64898/2026.07.31.742155

**Authors:** Vincent Guacci, Nicola Minchell, Tik Sung, Sergey Venev, Job Dekker, Douglas Koshland

**Affiliations:** Department of Molecular and Cell Biology, University of California, Berkeley, Berkeley, California 94720, USA; Program in Systems Biology, Department of Biochemistry and Molecular Pharmacology, University of Massachusetts Medical School, Worcester, MA, USA; Howard Hughes Medical Institute, Chevy Chase, MD, USA

## Abstract

The protein complex cohesin plays critical roles in genomic stability by tethering together sister chromatids at their pericentric regions and along their arms from S phase until anaphase. Cohesin-mediated pericentric cohesion prevents aneuploidy by ensuring bipolar attachment of sister kinetochores. Arm cohesion prevents loss of heterozygosity by biasing DNA repair via recombination between sister chromatids rather than between homologs. Here, we investigate in yeast whether cohesin also enhances genomic stability by suppressing unequal sister chromatid exchange (USCE) between repetitive sequences. In wild-type cells, the USCE rate between repeats 4kb apart (proximal) was 15X higher than repeats 68kb apart (distal). The USCE between distal repeats but not proximal repeats increased 4 to 7-fold in mutants with altered cohesin subunits or auxiliary factors. The level of increased distal USCE corresponded with reduced arm cohesion, reduced density of cohesion arm sites, and higher sister loci mobility. Our results suggest that high density of arm cohesion sites confines repair of DNA damage to local sequences. When the density of cohesion sites decreases, sister chromatid sequences are less confined, thereby enhancing distal repeat interactions and USCE. Another set of mutations disrupted both DNA replication and cohesin loading at the replication fork during S phase. Remarkably, distal USCE in these mutants increased approximately 100-fold and was 6-fold more likely than proximal USCE. This preferential hyperdistal USCE can be explained by an aberrant sister-chromatid structure that is normally prevented by proper coupling of cohesin function and replication.

## Introduction

Cohesin mediates higher-order chromosome structure in eukaryotes by its ability to tether DNA sequences within and between sister chromatids and to form chromosomal loops by extruding DNA. (1–5). These activities enable cohesin to perform three biological functions critical for genome stability. First, cohesin tethers together sister chromatids at the centromere and along the chromosome arms from the time of their synthesis in S phase until their segregation in mitosis (2, 6). This sister chromatid cohesion around the kinetochore is essential for biorientation of the sister chromatids on the mitotic spindle and for their proper segregation during anaphase (7, 8). Second, sister chromatid cohesion along the arms reduces the uncovering of deleterious recessive traits by mitotic recombination, termed loss of heterozygosity (LOH). A double-strand break (DSB) in a unique sequence on one sister chromatid is preferentially repaired by recombination using the spatially proximal identical sequence on the cohesed sister chromatid rather than a distal homolog (9, 10). This bias leads to the genetically neutral outcome of sister chromatid exchange (SCE) rather than potential LOH from mitotic recombination. Third, a recent study suggests that cohesin’s looping activity promotes efficient repair of a broken chromosome by acting locally at the site of a DNA double-strand break to direct the search for homologous repair sequences on the unbroken sister chromatid (11).

Here, we investigate an additional function of cohesin in genome stability, its ability to limit unequal sister chromatid exchange (USCE). USCE can occur and be problematic because repetitive DNA sequences are widespread in most genomes. When a DSB occurs in a repetitive element on one sister chromatid, the repair of the break can occur by recombination using an undamaged repeat on the sister chromatid, either at the homologous position (SCE) or at an ectopic position on the sister (henceforth abbreviated as the homologous or ectopic repeat, respectively). SCE leaves the sequences of both sisters unchanged. In contrast, unequal sister chromati exchange (USCE) between the damaged and ectopic repeats results in the duplication of the intervening sequence between the repeats on one sister chromatid and deletion of the intervening sequence between the repeats on the other sister chromatid. USCE-induced duplications and deletions will be small when the recombining repeats are close (proximal repeats), and large if repeats are far apart (distal repeats).

Cohesin’s role in limiting USCE has not been examined. Cohesin is a conserved protein complex composed of a heterodimer of the Smc1 and Smc3 sub-units along with the Mcd1/Scc1/Rad21 subunit and Scc3/Stag subunit (2, 12). Cohesin is enriched at chromosomal sites distributed along chromosome arms. In budding yeast, these enriched sites are termed cohesin-associated regions (CARs) and occur at ∼15kb intervals (13–15). Cohesin’s association with DNA and its looping activity depends on a loader complex and cohesin’s ATPase activity (4, 5, 16–18). In yeast, the Eco1p acetyltransferase and cohesin are recruited to the replication fork by associating with the Ctf4, Ctf18, and Chl1 proteins (19–21). Eco1p acetylates a subset of cohesins, **l**eading to the downregulation of their ATPase activity (18, 22, 23), the promotion of cohesion (24–26), and the suppression of looping (27–29). Cohesin acetylation also inhibits Wpl1p (25), a cohesin regulator that removes cohesin from chromosomes (30, 31). Eco1p acetylation of the K113 residue of the Smc3 subunit is essential for the establishment of cohesion (24–26).

We and others have identified mutations in yeast through loss-of-function and suppressor screens (Table 1) that alter cohesin’s ATPase activity, acetylation, chromosome binding, and cohesive activities (18, 23, 25, 32). Here, we exploit these mutations to test whether cohesin promotes genomic stability by limiting USCE in yeast and, if so, which of cohesin’s biological and biochemical activities are required for this function.

**Table 1.**
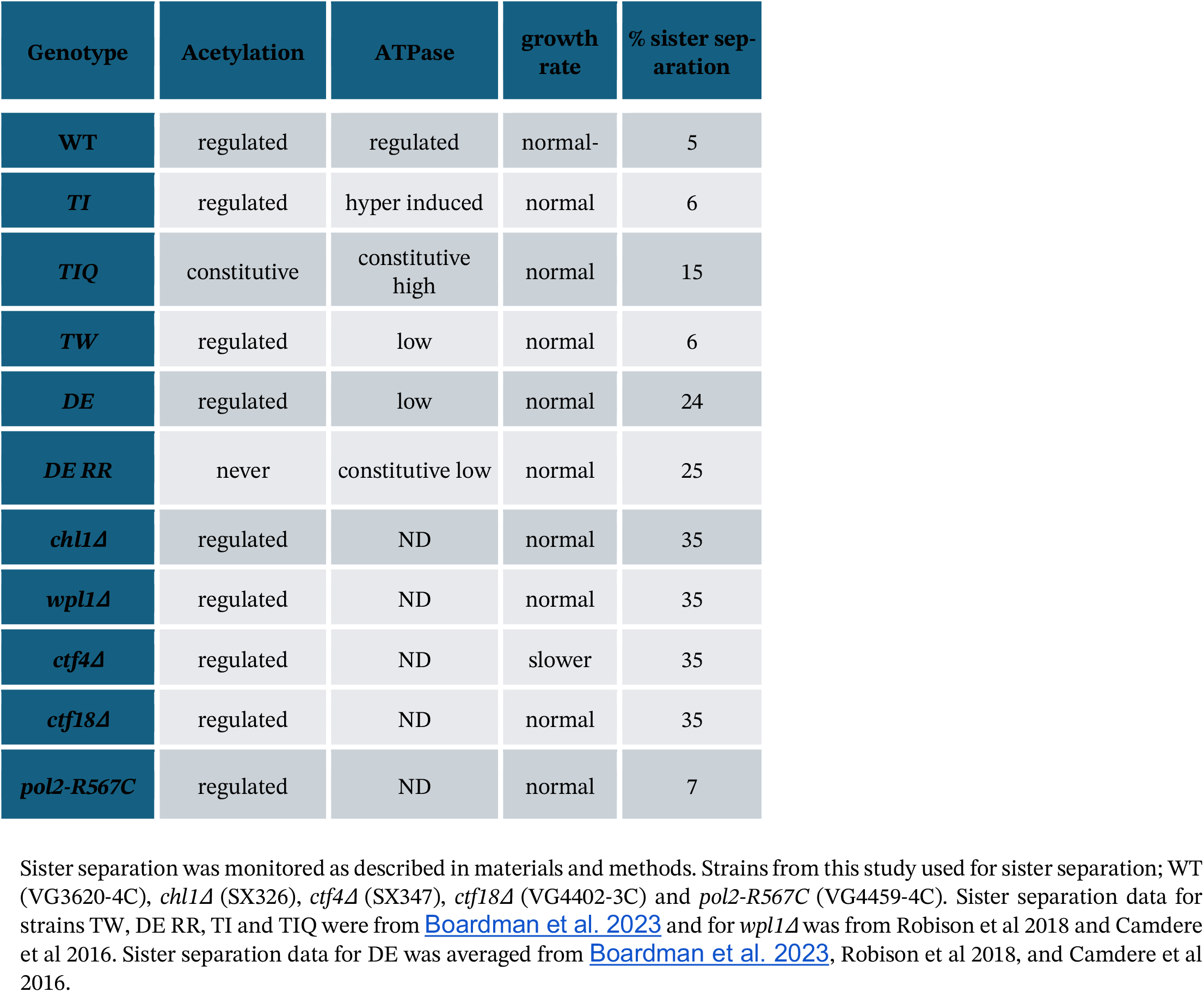

| Genotype | Acetylation | ATPase | growth rate | % sister separation |
| --- | --- | --- | --- | --- |
| **WT** | regulated | regulated | normal- | 5 |
| **TI** | regulated | hyper induced | normal | 6 |
| **TIQ** | constitutive | constitutive high | normal | 15 |
| **TW** | regulated | low | normal | 6 |
| **DE** | regulated | low | normal | 24 |
| **DE RR** | never | constitutive low | normal | 25 |
| ***chl1Δ*** | regulated | ND | normal | 35 |
| ***wpl1Δ*** | regulated | ND | normal | 35 |
| ***ctf4Δ*** | regulated | ND | slower | 35 |
| ***ctf18Δ*** | regulated | ND | normal | 35 |
| ***pol2-R567C*** | regulated | ND | normal | 7 |

Sister separation was monitored as described in materials and methods. Strains from this study used for sister separation; WT (VG3620-4C), *chl1Δ* (SX326), *ctf4Δ* (SX347), *ctf18Δ* (VG4402-3C) and *pol2-R567C* (VG4459-4C). Sister separation data for strains TW, DE RR, TI and TIQ were from Boardman et al. 2023 and for *wpl1Δ* was from Robison et al 2018 and Camdere et al 2016. Sister separation data for DE was averaged from Boardman et al. 2023, Robison et al 2018, and Camdere et al 2016.

## Results

### Increasing the distance between repeat sequences on a chromosome dramatically decreases USCE

To begin our interrogation of cohesin’s role in limiting USCE, we modified a previously described genetic assay for USCE in yeast (33). We generated 5’ and 3’ fragments of the hygromycin resistance gene, 5’ hyg and 3’ hyg, respectively. These hyg fragments share a 569 bp sequence from the middle of the hygromycin open reading frame (Fig. 1A). The fragments were integrated in the middle of the same chromosome arm (Fig. 1B top). The truncated fragments did not confer hygromycin resistance. However, an USCE using the shared 569 bp homology generates a sister chromatid with a full-length hygromycin gene (HYG) and a duplication of the chromosomal sequence between the original repeats (Fig. 1B middle). The other sister chromatid suffers a deletion of the chromosomal sequence between the original repeats (USCE by reciprocal exchange) so lacks a functional hyg gene. The cell inheriting the sister chromatid with the full length HYG gene is resistant to hygromycin (Fig. 1B, bottom right). We determined the frequencies of hygromycin-resistant descendants in 10 independent colonies derived from a hygromycin-sensitive founder cell. Using the method of the median (34), these frequencies were converted to a rate of hygromycin resistance generation, a direct measure of the rate of USCE (Materials and Methods).

**Figure 1.**
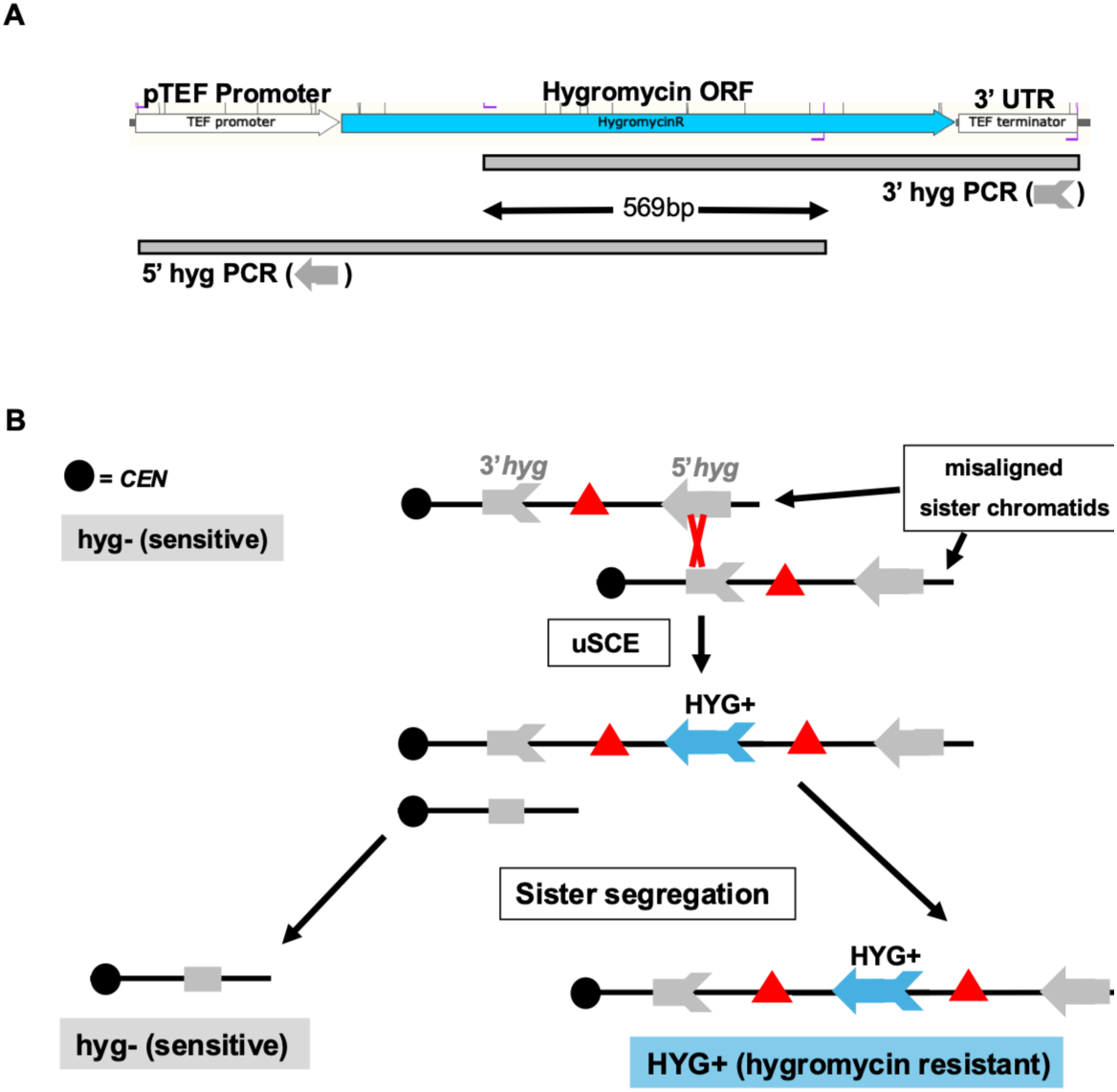
Schematic for assessing unequal sister chromatid exchange (uSCE) using a split hygromycin gene (HYG). A) Overlapping 5’hyg and 3’hyg fragments of the Hygromycin gene were made by PCR (grey bars). HYG orf (blue), pTEF promoter (white, left side) and 3’UTR (white, right side). The 5’hyg fragment contains the promotor and N-terminal 2/3 of HYG orf whereas and the 3’hyg fragment contains the C-terminal 2/3 of HYG orf and the terminator. Black arrows lines mark the 569bp orf sequence shared by the 3’hyg & 5’hyg fragments. B) Cartoon illustrating USCE between 3’hyg and 5’hyg. The 3’hyg (gray arrowtail) and 5’hyg (gray arrowhead) fragments are integrated at different sites along a chromosome. Sister chromatid misalignment and crossover in the region common to both fragments (USCE) generates a full length hygromycin gene (HYG+, blue arrow) on one sister chromatid along with duplication of genes between the hyg fragments (red triangles). The other sister chromatid contains only the shared hyg region (grey box) and is deleted for genes between the fragments. When sister chromatids segregate at anaphase, one daughter cell is hygromycin resistant (HYG+, right chromatid) whereas the other daughter cell remains hygromycin sensitive (left chromatid).

We constructed four reporter strains to measure the proximal and distal USCE rates on two different chromosomes, V and X as follows. First, we integrated a 3’ fragment of the hygromycin resistance gene (HYG) in the middle of an arm of either chromosome V or chromosome X in wild-type haploid strains (Suppl Fig S1A and S1B). For the two proximal USCE reporter strains, we then integrated a 5’ hyg fragment 4 kb away from the 3’ hyg fragment on the same chromatid. For the two distal reporter strains, the 5’ hyg fragment was integrated 68 kb away from the 3’hyg fragment. The hyg fragments were integrated in noncoding regions far from centromeres or telomeres to eliminate potential confounding factors associated with gene inactivation or specialized chromosome structures.

We used our reporter strains to determine the rate of hygromycin resistance/USCE as described above. For chromosome V, the proximal repeat USCE rate (7.9 × 10^-6^) was 15.1 times higher than the distal repeat USCE rate (0.52 × 10^-6^) (Fig. 2A & 2C). In our chromosome X reporter strains, the proximal repeat USCE rate (5.7 × 10^-6^) was 20.1 times higher than the distal repeat USCE rate (0.29 × 10^-6^) (Fig. 2B & 2C). Similar values for proximal repeat USCE and a decreased mitotic recombination rate as a function of distance have been reported previously (9, 33). (Lichten & Haber Genetics [1989] 123:261-268). The proximal USCE rates on chromosome V and X were very similar, and the distal repeat USCE rates on both chromosomes were also similar. These results suggested that the proximal and distal repeat USCE rates were governed by shared features beyond primary DNA sequence, and that one or more of these features were responsible for the distal USCE rate being so much lower than the proximal USCE rate.

**Figure 2.**
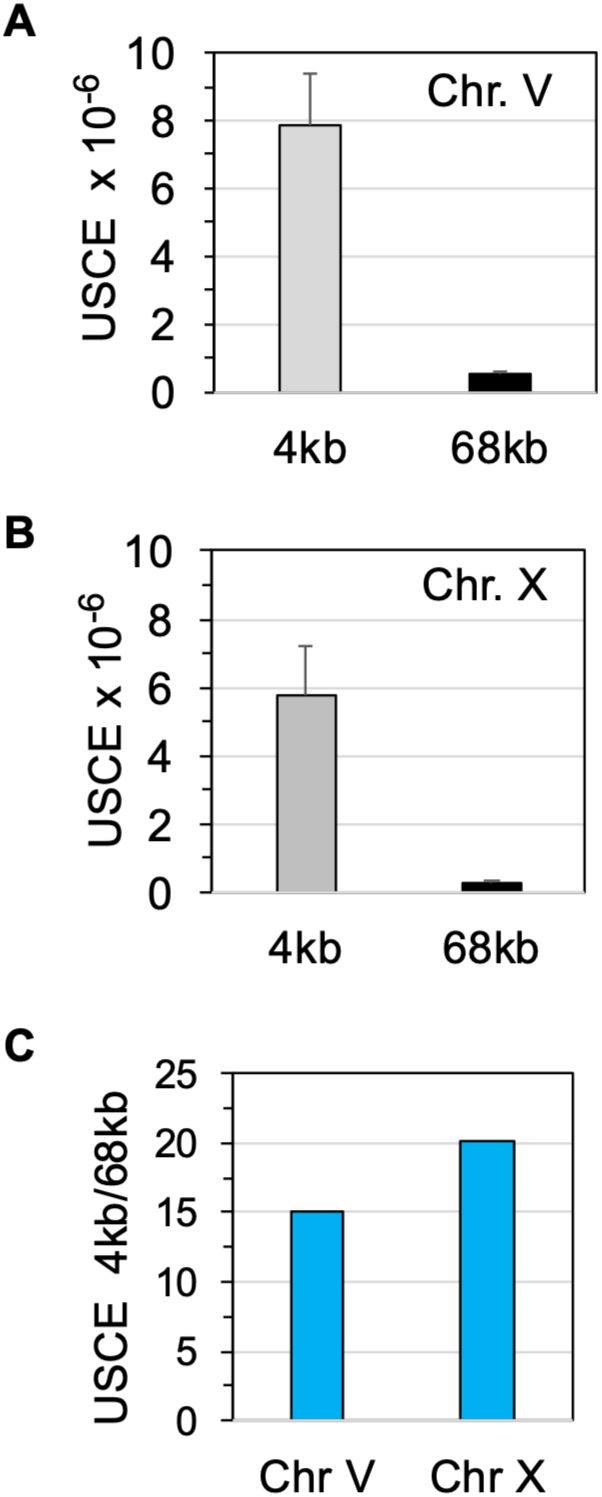
USCE rate is reduced as a function of distance. Comparison of proximal and distal USCE rates on chromosomes V and X. Proximal USCE rates were measured in wildtype haploid strains bearing 3’hyg and 5’ hyg fragments separated by 4kb whereas distal USCE rates determined using fragments separated by 68kb as described in Materials and Methods. A) USCE rates on chromosome V. Proximal USCE (4kb) and distal USCE (68kb) rates were determined using VG4220-14B and VG4222-12D, respectively. Data is from at least 4 independent experiments. B) USCE rates on chromosome X. Proximal USCE (4kb) and distal USCE (68kb) rates were determined using 4460-13C and 4436-1A, respectively. Data is from 2 independent experiments. C) The ratio of proximal to distal USCE rates was determined by dividing proximal rate by distal rate using data from A & B.

### Cohesin limits USCE between distal but not proximal repeats

We hypothesized that distal USCE could be limited by the higher-order structure of sister chromatids imposed by cohesin’s tethering or looping activities. If so, mutations that reduce cohesin function might impact USCE of distal repeats. Cohesion between sister loci is one sensitive biological readout of their impact on cohesin functions. In yeast, sister chromatid cohesion is assayed by FISH or by marking loci with LacO arrays that bind LacI-GFP (35, 36). In wild-type cells, sister chromatid arms are held in such close proximity that monitored loci appear as a single FISH signal or LacI-GFP signal in almost all cells. In cohesin-null mutants, sisters separate so that 70-90% of cells exhibit two signals (7, 8, 37). We exploited a panel of mutations that caused the separation of sisters at the *LYS4* locus, ranging from 6% to 35% (Table 1).

All the mutations increased the distal USCE rates on chr. V (Fig. 3A). Moreover, the fold increase in the distal repeat USCE rates in the mutants relative to wild-type correlated well with the increased level of sister separation at the *LYS4* locus (Fig. 3B). To determine whether the impact of cohesion dysfunction on USCE was sequence- or chromosome-specific, we introduced a subset of these mutations into our chr. X distal USCE reporter strain. The increase in the USCE rate and the fold change compared to wild-type were remarkably similar to those seen in the chr. V distal USCE reporter strain (Suppl Fig 2A). These results showed that cohesin function helped limit distal USCE, and the extent of this repression depended on the level of cohesin function. Additionally, cohesin’s suppression of distal USCE was not chromosome-specific.

**Figure 3.**
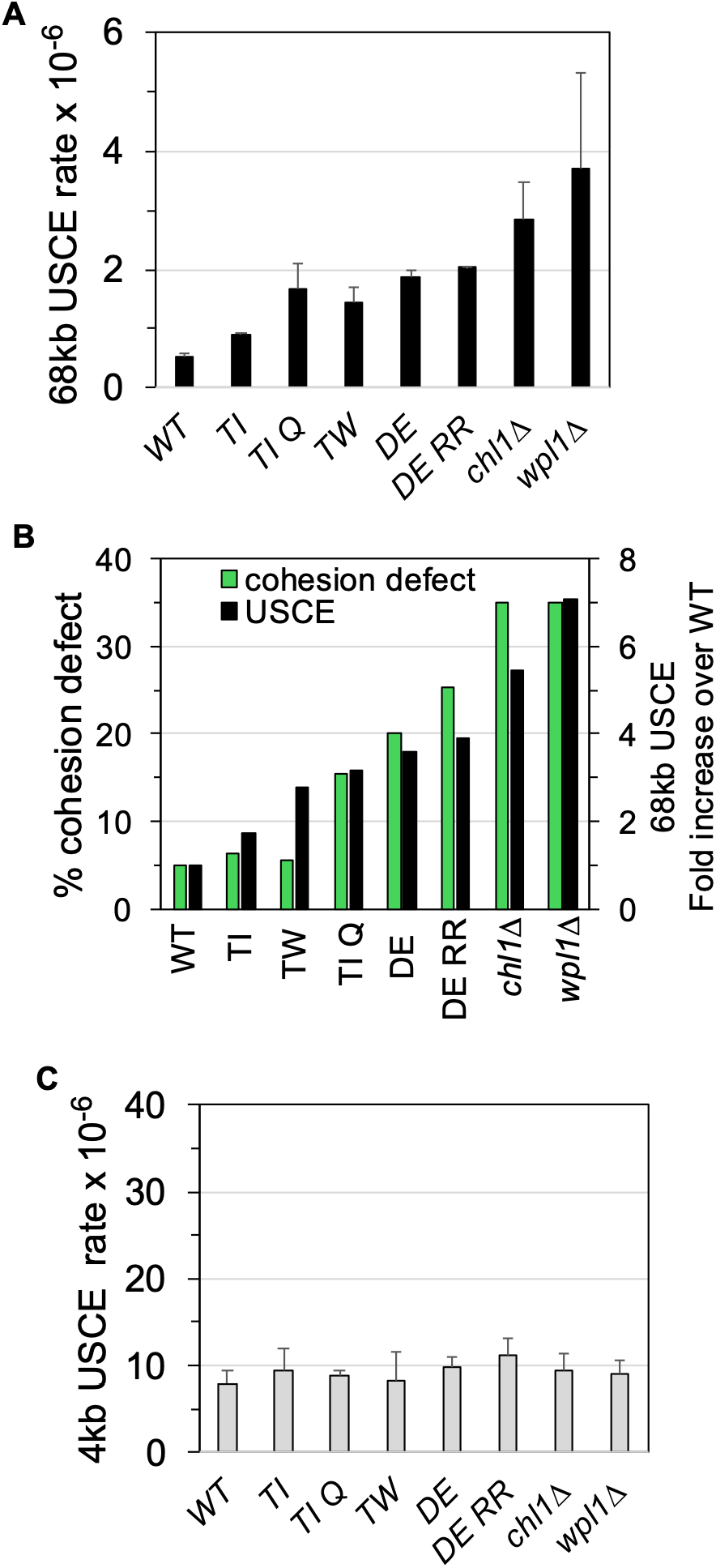
Cohesin mutants increase distal USCE but have no effect on proximal USCE. A) Effect of cohesin mutants on distal USCE rates. Cohesin mutants were introduced into wildtype strain VG4222-12D [WT] to form mutant haploids VG4236-6B (*smc1-T117I* [TI]), VG4260-15D (*smc1-T1117I smc3-K113Q* [TIQ], VG4240-10B (*smc1-T1117W* [TW]), VG4232-8C (*smc1-D1164E* [DE]), VG4258-5D (*smc1-D1164E smc3-K112R, K113R* [DERR]), VG4250-13D [*chl1Δ*] and VG4228-4B [*wpl1Δ*] then strains assayed for distal USCE rates. Data is from 2 to 3 independent experiments. B) The increase in distal USCE correlated with the level of the defect in sister chromatid cohesion. The fold increase of distal USCE in each mutant was calculated by dividing the rate of distal USCE in mutants by the rate for WT derived in (A). The percentage cohesion defect at the *LYS4* locus for each strain (see Table 1) was plotted on the same graph as USCE. C) Effect of cohesin mutants on proximal USCE rates. Cohesin mutants were introduced into wildtype strain VG4220-14B [WT] to form mutant haploids VG4235-5A (*smc1-T117I* [TI]), VG4259-13B (*smc1-T1117I smc3-K113Q* [TIQ], VG4239-9C (*smc1-T1117W* [TW]), VG4231-7A (*smc1-D1164E* [DE]), VG4275-3B (*smc1-D1164E smc3-K112R, K113R* [DERR]), VG4249-7A [*chl1Δ*] and VG4227-3B [*wpl1Δ*

To assess whether cohesin mutants affected proximal USCE, we introduced the mutation panel into our proximal USCE reporter strains. The USCE rates for all mutants were similar to the wild-type for both chromosome V and X (Fig. 3C & Suppl Fig 2B). Therefore, in contrast to distal repeats, the USCE for repeats separated by 4 kb was not limited by cohesin function. This observation rules out an alternative explanation whereby the increased distal USCE seen in cohesin mutants was merely due to increased DNA damage. If this were the case, the increased damage should have also elevated proximal USCE. Alternatively, wild-type cohesin creates a structure(s) or organization of sister chromatids that inhibits distal repeats interactions. Cohesin mutants disrupt this structure/organization, which enables distal repeats to interact and recombine at higher rates. If so, this putative inhibitory structure must function at a scale larger than 4 kb since the proximal USCE rate was not affected by cohesin mutants.

The increased USCE rates in the mutants prompted the question whether these increases correlated with specific perturbations of cohesin activities or regulation. In our mutant panel the DE (*smc1D1164E*) and TW (*smc1-T1117W*) mutations reduced cohesin ATPase activity while the TI (*smc1T1117I*) and TIQ (*smc1-T1117I smc3-K113Q*) mutations increased it (18, 23). Three mutations alter cohesin’s response to acetylation. The DE and TW mutations allow cells to form sister chromatid cohesion and survive in the absence of K113 acetylation (18, 23). In contrast, the TIQ mutation allows cells to live and form cohesion when K113 is mutated to K113Q, mimicking a constitutively acetylated state (23). Finally, two mutations alter the regulation of cohesin binding to DNA by either reducing its loading on DNA during DNA replication (*chl1Δ*) or increasing its residence time (*wpl1Δ*) (21, 38). Distal USCE rate was elevated to the same level in mutants that elevate or reduce ATPase activity (TIQ and DE) respectively, eliminate acetylation or constitutively mimic acetylation (DE RR and TIQ), or alter cohesin loading or residence time (*wpl1Δ* and *chl1Δ*) (Fig. 3A), indicating that compromising different aspects of cohesin function led to similar increases in distal USCE rate. Furthermore, our results suggested that presence of ATPase activity or cohesin acetylation were not sufficient to suppress USCE. Rather, maintaining proper regulation of cohesin acetylation and ATPase levels were important for suppressing distal repeat USCE.

### Distal repeats are less likely to interact than proximal repeats

An explanation for the decreased distal repeat USCE rate in wild-type cells was that chromosome structure limited interaction of distal repeats more than proximal repeats. Using our previously published Micro-C dataset (39), we were able to assess the interactions between the chromosomal sequences where the 3’ and 5’ *hyg* repeats were inserted in our USCE reporter strains. Robust interactions were observed between CARs flanking the proximal repeat insertion sites and among intervening sequences between the CARs on chr. V and chr. X (Suppl Figs 1A & 1B, upper left oval). In contrast, such CAR-CAR and intervening sequence interactions were undetectable between distal repeats (Suppl Figs 1A & 1B, upper right oval). These results support the hypothesis that the reduced distal USCE rates in wild type cells was due to a lower likelihood of interactions between distal repeats than between proximal repeats. In contrast, in the *wpl1Δ* mutant, interactions were detected between distal chromosomal sequences and distal CARs (Suppl Figs 1C & 1D, upper right oval). The *wpl1Δ* mutant had the highest increase in distal USCE. Thus, the increased ability of distal sequences to interact in *wpl1Δ* cells provides a mechanism to explain why suppression of distal USCE is weaker than in wild-type cells. An increased probability of interaction leads to higher distal USCE.

### Suppression of distal USCE correlates with a higher density of cohesion sites along chromosome arms, which potentially limits both loop size and sister loci mobility

We reasoned that the presence of cohesion sites between sisters could reduce distal repeat interactions. ChIP of cohesin subunits identified genomic sites of stable cohesin binding, called CARs, which occur approximately every 15 kb along chromosomal arms (14, 15, 40). CARs have been assumed to be likely sites of arm cohesion. The 68 kb genomic regions between the *hyg* distal repeats on chromosomes V and X contain multiple CARs, suggesting at least one or more cohesion sites existed between them. In contrast, the proximal *hyg* repeats were positioned between two CARs, and hence the proximal repeats were likely flanked by, rather than separated by, cohesion sites.

We used sisterC to estimate the likelihood of a CAR site being a cohesion site as previously described (41). This chromosome conformation method allows one to distinguish whether a DNA site on a chromatid is interacting with DNA sequences from the same chromatid (intra) or with its sister chromatid (inter). In wild-type, the probabilities for intra- and inter-sister chromatid interactions both decreased as a function of the chromosomal distance between the sequences. At short distances, the interaction probabilities for pairs of intra-chromatid sequences were higher than inter sister chromatid sequences, but the probabilities converged around 25 kb distance (Fig. 4A and Suppl Fig 4A), consistent with previous results (41). The convergence should mark the distances between cohesion sites because the close proximity of sister sequences at cohesion sites makes these DNA loci equally likely to interact with distal sequences on the same chromatid or on the sister chromatid. This 25 kb cohesion spacing, coupled with CAR spacing of ∼15 kb, suggested that on average every other CAR is a site of cohesion. From this periodicity of CARs as cohesion sites, we concluded that the 68 kb intervening sequences between the distal *hyg* repeats were likely to have multiple cohesion sites. In contrast, the 4 kb spacing between proximal repeats are not separated by any CARs, and consequently, no cohesion sites. Thus, the increased likelihood of a cohesion site(s) being present between the distal repeats is consistent with the idea that cohesion sites contribute to suppression of distal USCE.

**Figure 4.**
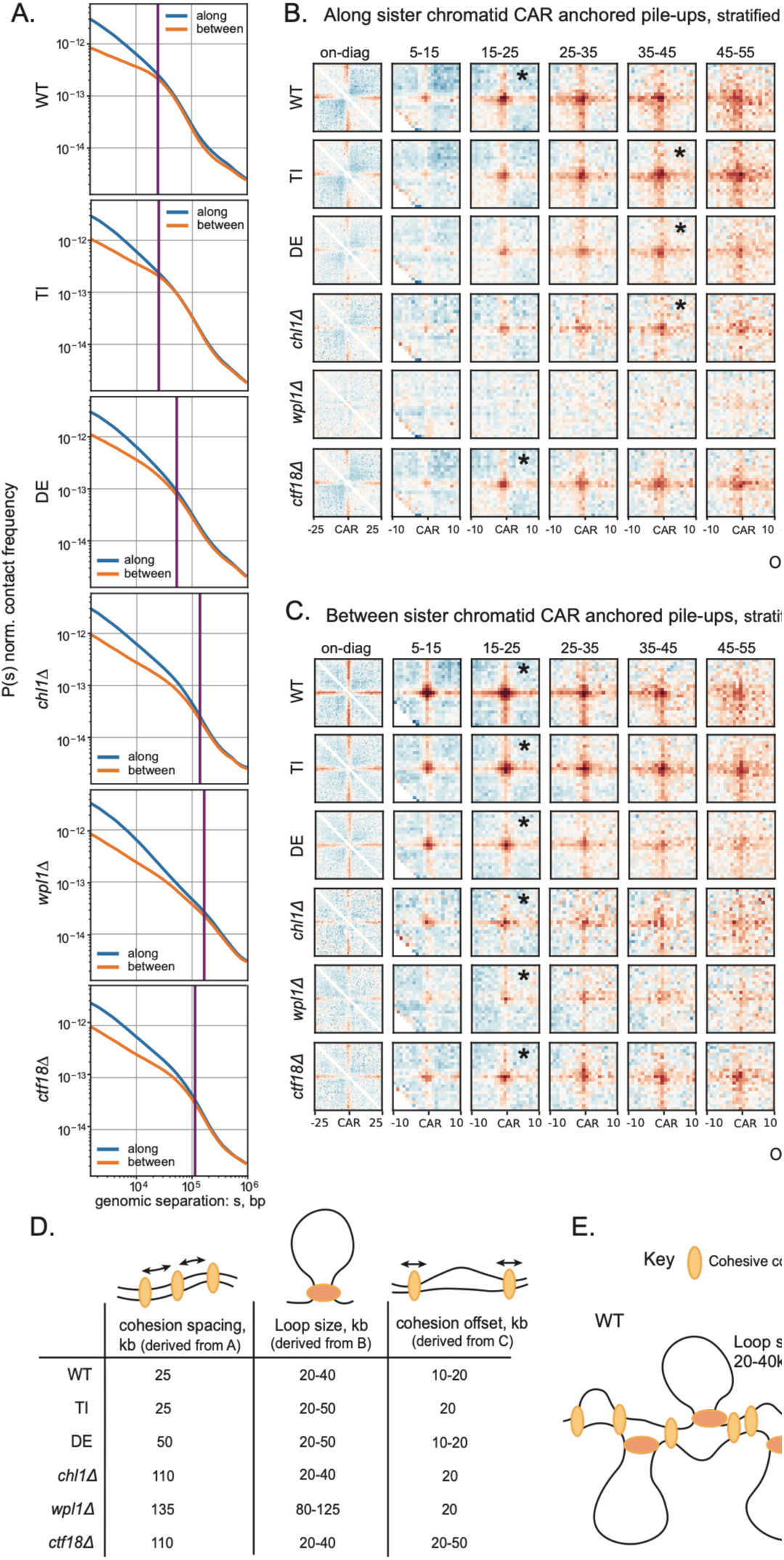
SisterC reveals the effects of cohesin mutants on chromatin interactions along and between sister chromatids. Sister-C was performed on six S.*cerevisiae* strains in YLV11 background. These are the parent wild-type YLV11 [WT], cohesin mutants affecting ATPase activity VG4308-4D (*smc1*- *D1164E* [DE]) and VG4322-5C (*smc1-T1117I* [TI]) and VG4312-1B [*chl1Δ*], VG4313-10C [*wpl1Δ*] and VG4309-6A [*cft18Δ*] mutants. Cells were synchronised to mitosis using nocodazole after one round of replication with BrdU. Two replicates are combined. (A) Contact frequency *P* plotted as a function of genomic separation for interactions either along or between sister chromatids. (B) Intra-sister [along] (C) Inter-sister [between] Pile-up interaction maps based on the distance between *WT* CAR’s (excluding peri-centromeric peaks), plotted at a resolution of 1kb. The asterisks indicate the maximum strength pile-up for each strain. A table deriving the distance between cohesive cohesins [cohesion spacing] from (A) (the intersection of the lines), the most frequent size of intra-sister loops [Loop size] from (B), and the most frequent offset of DNA sequence between two sisters [cohesion offset] from (C) for all strains. Most frequent loop size and cohesion offset were manually chosen based on strength from the quantification of the pile-ups for each strain (see Supp Figure 4). (E) Schematic representing the effects on sister chromatids of cohesive and extruding cohesin in wt cells.

To test whether genome-wide cohesion site spacing was altered in our mutants, we introduced a subset of mutations (*DE, wpl1Δ, chl1Δ* and *TI*) into the sister-C strain background. We then performed sisterC to determine the genomic distance at which the probabilities of intraand inter-sister chromatid sequence interactions converged, marking cohesion sites. This convergence occurred at greater distances in all the mutants except TI, suggesting that these mutations caused increased cohesion spacing with *wpl1Δ* being the most severe (Fig. 4A and Suppl Fig 4A). The increasing cohesion spacing in the mutants correlated extremely well with a higher distal repeat USCE rate (Suppl Fig 4B). This result aligns with the model that a higher density of cohesion sites on chromosomal arms suppresses distal USCE by limiting the interaction of repeats. Interestingly, the *wpl1Δ* mutant also exhibited increased trans cen-cen interactions and larger loops emanating from the centromeres (Supp Fig S3), two traits that did not correlate with elevated distal USCE in the other mutants but may reveal additional important Wpl1 functions to pursue in further studies.

Our mutants could also have enhanced distal USCE through mechanisms that actively promoted the interaction of distal sequences such as loop extrusion (42–45). Larger loops could bring the distal sequences along and between sisters into closer proximity, thereby boosting chances for USCE. Alternatively, cohesin mutations could alter the choice of cohesion tethering sites on sister chromatids during cohesion establishment. Since cohesion is coupled to replication, the sites of tethering on the sister chromatids in wild-type cells occur at homologous positions or nearly homologous positions (cohesion offset). By perturbing the coupling of replication and cohesion establishment or by increasing loop size, cohesin mutations could increase the offset of cohesion, consequently increasing the probability of interactions between distal sequences on sister chromatids.

With these models in mind, we employed sisterC to measure loop size and cohesion offset in the mutants. The cohesion offset in the mutants varied in strength but all were able to maintain a wild type level of offset with some mutants having a wider range (Figs. 4B and 4D). However, the loop size in the mutants differed from wild type but did not correlate with the distal repeat USCE rate (Suppl Fig 5). Because none of the se mutations altered cohesion offset, they did not address a potential role of minimizing cohesion offset in limiting distal USCE.

### USCE suppression correlates with constraining independent sister chromatid mobility

Multiple cohesion sites between the distal repeats could limit their interactions by constraining their independent mobility. To test this possibility, we used the microtubule inhibitor nocodazole to mitotically arrest yeast cells which contain a GFP-tagged *LYS4* arm locus. We imaged GFP every 20 seconds for almost 7 minutes (Movie S1 and S2). In wild-type cells, approximately 10% of cells exhibited two spots at any given time (columns) and that over the time course (rows), two spots were rarely observed in most individual cells (Fig. 5). These results suggested that most wild-type cells had sufficient cohesion sites proximal to the *LYS4* locus arms to restrict the mobility of the marked sister loci, thereby keeping them close together except for rare transient separation events. A small subset of cells had a much higher probability of having two spots over time than predicted from the fraction of cells in the population with two spots, suggesting these cells had a reduced probability of arm cohesion sites near the *LYS4* arm locus. This reduction could be due to a stochastic process naturally governing cohesion site establishment, or because a few cells escaped the metaphase arrest imposed by nocodazole and initiated sister chromatid dissolution. In contrast, for cells lacking cohesin function (*mcd1-AID*), almost all cells in the population had two spots at a given time, and almost all individual cells had two spots over the time course.

**Figure 5.**
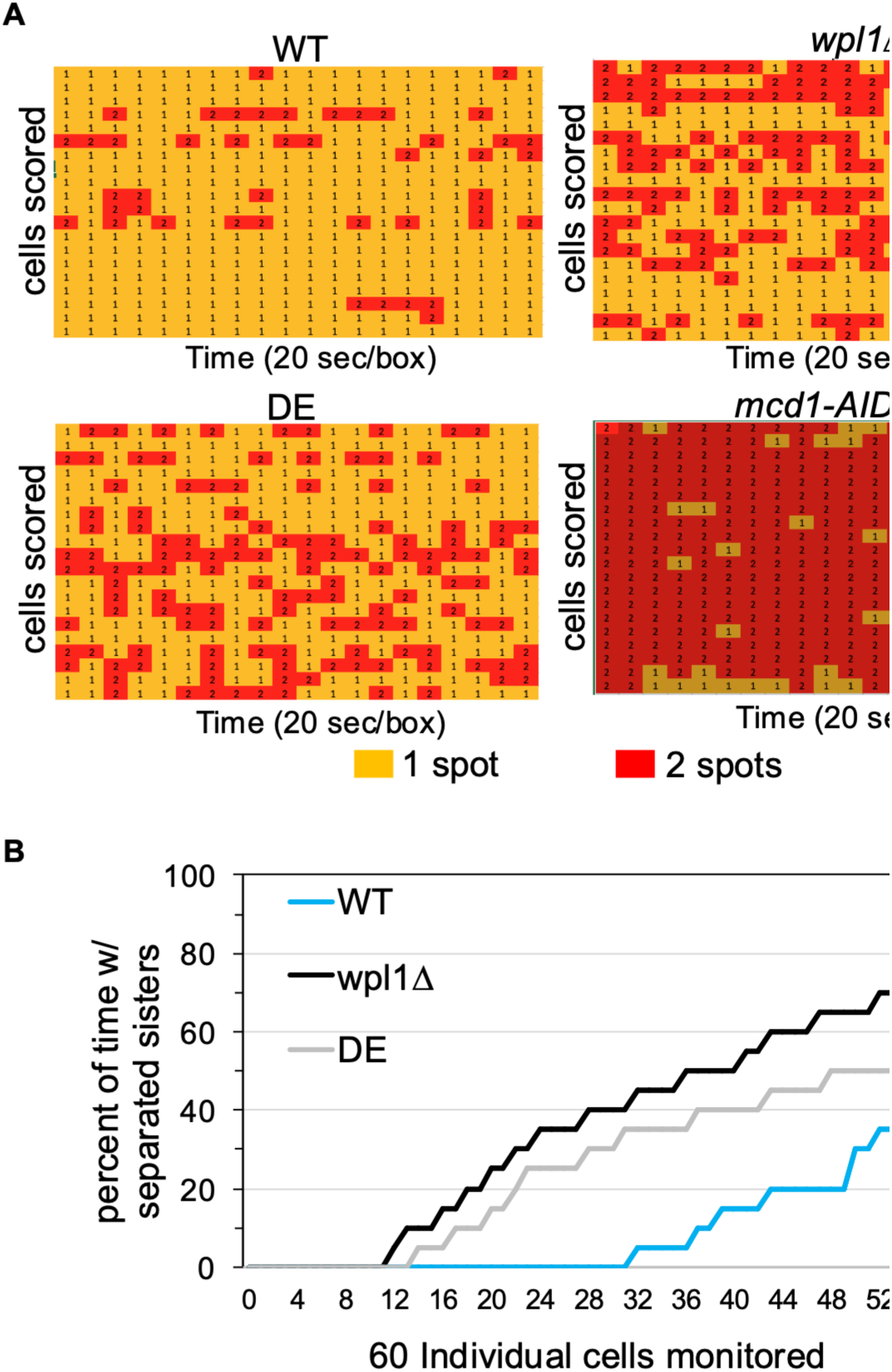
Live imaging to monitor sister separation in real time. Haploid wildtype strain VG3620-4C [WT] and cohesin mutant strains VG4138-2A (*smc1-D1164E* [DE]), VG3956-14A [*wpl1Δ*] and VG3902-3A [MCD1-AID] were assayed for cohesion loss at the *LYS4* locus. Cells were arrested in G1 then released into nocodazole to arrest in metaphase and processed for live cell imaging (Materials and Methods). Auxin was added to the *MCD1-AID* strain from G1 phase through metaphase phase. (A) For each strain, live cell imagining of 20 cells was performed every 20 seconds for 400 seconds and scored for whether the *LYS4* locus had sister chromatid cohesion (1 spot; yellow box) or had lost cohesion (2 spots, red box). Shown is data from one of 3 independent experiments. (B) Data from live cell imaging of 60 cells of WT, DE and *wpl1Δ* from 3 experiments plotted showing the percentage of time each cell exhibited sister separation. The delay in the curve start reflects the number of cells which never exhibit sister separation, which is greatest for WT as more than half these cells never exhibit any separation.

In the *DE* and *wpl1Δ* mutants, the fraction of time sister loci were separated in individual cells increased dramatically compared to wild-type, and this increase correlated with distal repeat USCE rate and cohesion spacing (Fig. 5A and 5B). Still, some cells in both mutants had few or no times where sister separation occurred. These results are consistent with a lower probability of cohesion sites being located near the *LYS4* arm locus in most *DE* and *wpl1Δ* cells than in wild type cells. Thus, live imaging established a correlation between increased independent sister loci mobility, increased cohesion site spacing, and increased distal repeat USCE rate. Together these correlations supported the hypothesis that the increased likelihood of cohesin sites between repeats repressed distal USCE by limiting repeat mobility and hence their interaction.

### Hyper USCE in the cohesin auxiliary factors associated with the replication fork

Cohesion is established during or soon after replication (46, 47). At the DNA replication fork, Chl1 and Ctf4 proteins associate with each other and the lagging strand, whereas Ctf18 protein associates with the leading strand (19, 48–50). A substantial fraction (30-35%) of *chl1Δ, ctf4Δ*, and *ctf18Δ* mutant cells arrested in metaphase exhibit separated sister loci (Table 1, (51, 52). Together, these data suggest that these factors help promote the establishment of cohesion, and their absence leads to a partial loss of arm cohesion. However, while *ctf4Δ* and *ctf18Δ* mutants exhibited slowed DNA replication (48, 53, 54) and Suppl Fig 6, the *chl1Δ* mutants exhibit normal replication (21, 38). This difference suggests that Ctf4 and Ctf18 proteins uniquely contributed to proper DNA replication and cohesion establishment.

To determine whether this coupling of cohesion and replication is crucial for inhibiting USCE, we used our assay to measure USCE rates in *ctf4Δ* and *ctf18Δ* mutants. Remarkably, the distal repeat USCE rate for chromosome V increased by 77-fold in *ctf18Δ* cells and 121-fold in *ctf4Δ* cells (Fig. 6A). Similarly, the *ctf18Δ* mutant exhibited a significant increase in the distal USCE rate for chromosome X (Suppl Fig 2C). The 68 kb regions in the chromosome V and X reporters did not share any apparent genome characteristics, such as the number or positions of replication origins relative to the repeats. Thus, the extremely high USCE rate in these strains was unlikely to be due to an unknown effect of *ctf4Δ* and *ctf18Δ* on specific sequences. In contrast, the proximal repeat USCE rates in *ctf4Δ* and *ctf18Δ* mutants were comparable to those of wild-type (Fig. 6B and Suppl Fig 2D), corroborating a previous study of proximal USCE in *ctf4Δ* cells (55). The fact that proximal USCE is not elevated suggested that the hyper distal repeat USCE rates were unlikely due to elevated DNA damage in these mutants.

**Figure 6.**
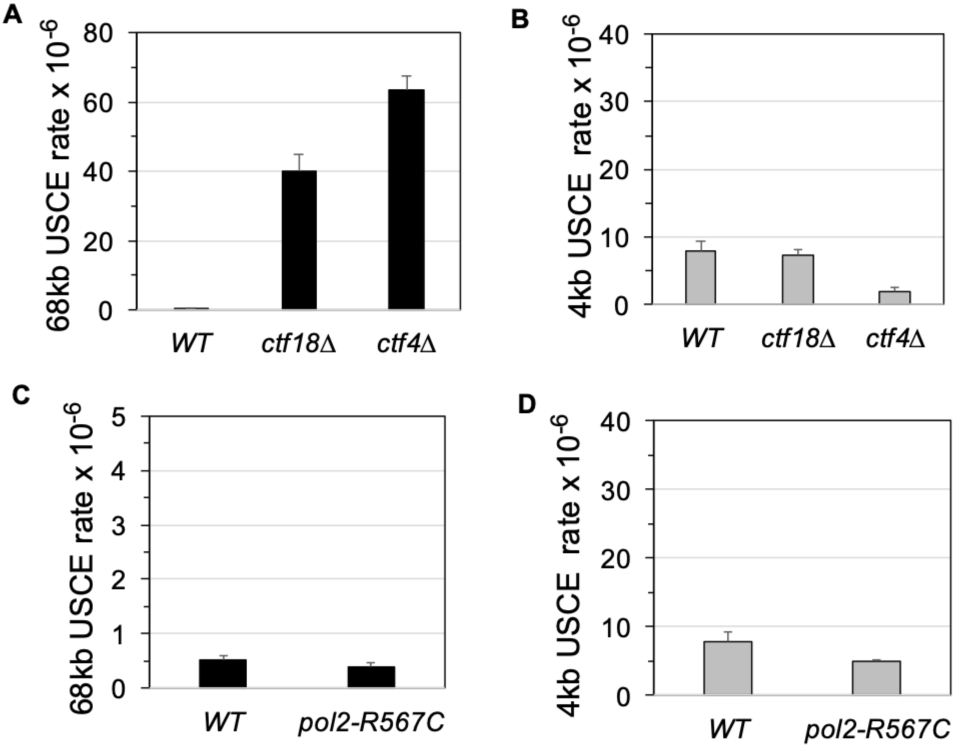
*ctf4* and *ctf18* mutants dramatically increase distal uSCE but do not increase proximal USCE. A & B) Effect of *ctf4* and *ctf18* mutants on USCE rates. A) Distal USCE rates. Wild-type haploid VG4222-12D (WT) and mutants VG4252-18D (*ctf18Δ*) and VG4281-1D (*ctf4Δ*) were assayed for USCE rates (Materials and Methods). B) Proximal USCE rates. Wild-type haploid VG4220-14B (WT) and mutant haploids VG4280-8B (*ctf4Δ*) and VG4251-10A (*ctf18Δ*) and assayed for rate of uSCE (Materials and Methods). C & D) Effect of *pol2-567C* on USCE rates. C) Distal USCE rates. Wild-type haploid VG4222-12D (WT) and mutant VG4425-9D (*pol2-R567C*) were assayed for USCE rates. D) Proximal USCE rates. Wild-type haploid VG4220-14B (WT) and mutant VG4423-4B (*pol2-R567C*) were assayed for USCE rates. Data for all strains in (A-D) were from 2 independent experiments.

We investigated the possibility that the increased distal repeat USCE rates in *ctf4Δ* and *ctf18Δ* mutants were solely due to their slow replication. For this purpose, we introduced the *pol2-R567* mutation into our distal and proximal USCE reporter strains. The *pol2R567* mutant also caused slow replication (56) but had wild-type levels of cohesion (Table 1). The USCE rates in both the distal and proximal repeats of the mutant were similar to wild-type, indicating that slowed replication alone did not elevate USCE rates (Fig. 6C & 6D). Consequently, the elevated distal repeat USCE rates in *ctf4Δ* and *ctf18Δ* were likely attributed to a change in the structure of the sister chromatids, possibly caused by the combination of slowed replication and a cohesion defect.

Two surprising features of the hyper distal repeat USCE rate in *ctf4Δ* and *ctf18Δ* mutants provide a clue to the cause. First, distal USCE rates were more than 10-fold higher than those in *wpl1Δ* and *chl1Δ* mutants (Fig. 3 and Fig. 6), despite exhibiting a similar increase in sister-loci separation. Second, the rate of distal repeat USCE was 5-8 higher than the rate of proximal repeat USCE in wild-type (Fig. 6A & 6B). Together, these surprising features suggested that the *ctf4Δ* and *ctf18Δ* mutations induced an additional change(s) to chromosome structure besides the loss of cohesion sites.

To further interrogate potential differences in chromosome structure in these mutants that could account for the hyper distal repeat USCE rate, we attempted to introduce both the *ctf18Δ* and *ctf4Δ* mutations into the strain background used for sisterC analysis. Only the *ctf18Δ* sisterC strain was able to be generated. The *ctf18Δ* mutant had increased cohesion spacing similar to the level seen in *wpl1Δ* and *chl1Δ* (Fig. 4A & 4D). The similar cohesion spacing correlated with the similar levels of separated sister *LYS4* loci seen in *ctf18Δ, ctf4Δ, chl1Δ* and *wpl1Δ* mutants (Table 1). It also reinforced the conclusion that the loss of cohesion sites in the *ctf18Δ* and *ctf4Δ* mutants was not solely responsible for the hyper-level of USCE rate in these mutants. The loop size in the *ctf18Δ* mutant was similar to that of the wild-type, suggesting that this parameter also did not contribute to the hyper-elevated USCE rate. Unlike all the other mutants, the cohesion offset increased in the *ctf18Δ* mutant compared to wild type (Fig 4D). These results suggested that coupling replication and cohesion establishment may suppress distal USCE possibly by limiting cohesion offset.

## Discussion

In this report, we describe an assay to assess unequal sister chromatid exchange (USCE) between repeats separated by 4 kb (proximal) and 68 kb (distal) on two yeast chromosomes. We use this assay to interrogate USCE in a panel of mutants compromised for cohesin’s ATPase activity (*DE, TI, TIQ*), acetylation regulation (*DE, TIQ*), recycling (*wpl1Δ*), and loading (*chl1Δ, ctf4Δ*, and *ctf18Δ*). In all these mutants, the USCE rate between distal repeats is elevated. These results demonstrate that cohesin helps to repress distal USCE. Cohesin could mediate this repression by preventing the formation of DNA damage that induces USCE or by altering the structure(s) of sister chromatids which normally hinders recombination between the distal repeats. If cohesin dysfunction increased DNA damage, then damage should also have induced USCE between proximal repeats, which did not occur in any of the mutants in our panel. Therefore, we conclude that sister chromatid structure or organization imposed by cohesin suppresses distal USCE, establishing another critical role for cohesin in genome stability beyond its established roles in chromosome segregation and DNA repair.

In this study, cohesin mutants fell into two classes, those exhibiting moderate increases in distal USCE rates and those exhibiting hyper distal USCE increases. Several observations from the moderate class suggest that the number of cohesion sites on the chromosome arms plays a key role in regulating distal USCE. Our ChIP-seq and sisterC analyses show in wild-type cells that the 68 kb intervening regions between distal repeats likely have multiple cohesion sites whereas the 4 kb regions between proximal repeats have none, correlating the intervening cohesion sites with lower distal USCE. Furthermore, we show that the distal USCE rate increases in mutants defective in cohesin or its associated factors, correlates with progressively greater sister locus separation, greater spacing between cohesion sites, and greater independent dynamic mobility of sister sequences. We conclude that the cohesion sites between distal repeats in wild-type cells suppress independent sister chromatid mobility, thereby inhibiting distal sequence interaction and distal USCE

Cohesin’s tethering and looping activities lead to two models for how cohesion sites could reduce independent sister sequence mobility and suppress USCE. The flanking cohesion sites could limit the diffusion of the proximal repair templates, making them preferentially found in an homology search.

Loss of cohesion sites flanking the damaged repeat would allow the proximal repeat on the sister to diffuse further away, thereby enhancing the distal repeat’s ability on the sister to compete. A second more intriguing model stems from two recent observations. Cohesin extrudes DNA at the site of damage to promote the homology search, and extrusion is blocked by cohesion sites (57–59). Thus, the presence of high density cohesion sites limits the homology search to domains between cohesion sites, preventing the search from finding distal repeats and inhibiting distal USCE. By reducing the cohesion sites, the mutations should extend the homology search by allowing bigger loops, and promote distal repeat USCE. While, we don’t see a strict correlation between increasing distance between cohesion sites and loop size in our mutants, it is important to remember loop size reported here is an average of loops genome wide and may not inform on the loop size associated with damage induced extrusion.

The *ctf4Δ* and *ctf18Δ* mutants exhibited hyper distal USCE with three unusual features. First, the distal repeat USCE rate in *ctf4Δ* and *ctf18Δ* mutants was 10 to 20-fold more than the strongest mutants in the moderate class (*wpl1Δ* and *chl1Δ*), despite having similar cohesion defects and cohesion site spacings. Second, in the *ctf4Δ* and *ctf18Δ* mutants, the distal USCE rate was 5-8 fold higher than the proximal rate while in the moderate mutant class, the distal rate never exceeded one third of the proximal rate. Third, despite these dramatic changes in the distal USCE, the proximal USCE rate was not altered. These unusual features of *ctf4Δ* and *ctf18Δ* mutants suggest that Ctf4p and Ctf18p suppress distal USCE by a distinct mechanism from the factors in the moderate USCE group. Furthermore, the fact that the two mutants share all these unusual USCE features including very similar levels of hyper distal USCE, suggest that their hyoerdistal USCE derives from a defect in a shared function.

Ctf4 and Ctf18 proteins both act at the DNA replication fork duing S phase to promotge proper DNA replication and cohesion establishment (51, 52); (48, 53, 54). We suggest that the common cause of the hyper-distal USCE in the *ctf4Δ* and *ctf18Δ* mutants is a failure to properly couple cohesin function to replication. This uncoupling leads to the formation of an aberrant sister chromatid(s) structure which is responsible for the preferential hyper distal USCE. A potential candidate for this aberrant structure came from our observation that *ctf18Δ* mutants increased cohesion offset. Increased offset would impact USCE in two ways (Fig. S5). First, it would bring distal repeats in closer proximity so that a damaged repeat would have a higher probability of interacting with and recombining with a distal repeat on the sister chromatid. Second, large cohesion offset would also move the damaged repeat away from homologous or any proximal repeats on the sister chromatid, reducing their ability to compete with a distal repeat as a repair template. However, the difference in the cohesion offset between *ctf18Δ* and the other moderate mutants like *wpl1Δ* may not be sufficient to explain the preferential hyper distal USCE, suggesting it may be caused by another yet undetected aberrant structure. Further studies to elucidate the mechanism of hyper distal USCE are critical as they will likely provide important insights into the biological significance of the coupling of cohesion establishment to the leading and lagging strands of the replication fork by Ctf18 and Ctf4 respectively.

In principle other shared defects of *ctf4* and *ctf18* mutants could cause the preferential hyper-distal USCE. Both mutants have elevated DNA damage and/or be defective in DNA repair (20, 60–63). Increasing DNA damage genome wide could elevate distal USCE. However, elevated DNA damage should have also stimulated proximal repeat USCE which was not observed in either mutant. Alternatively the *ctf4Δ* and *ctf18Δ* mutations could have elevated damage specifically at the distal repeat because of some mutation-sensitive feature of the surrounding chromatin. However, no such localized damage in these mutants has been reported. In addition, since we observed hyper distal USCE on both chromosomes V and X, this putative sequence specific chromatin feature would have to have been fortuitously present at the sites of integration of the distal repeats on both chromosomes. The c*tf4Δ* and *ctf18Δ* mutants also exhibit reduced acetylation, cohesin binding to CARs and slow replication (64); (48, 53, 54). However, *chl1Δ* also reduced cohesin acetylation and cohesin binding at CARs (21, 38) but did not exhibit the hyper-elevated distal repeat USCE (this study). In addition, we found that the *pol2-R567C* mutation, which causes slow DNA synthesis, did not elevate distal repeat USCE rate nor does it exhibit a cohesion defect. Together, these observations show that elevated general or sequence specific DNA damage, the loss of cohesion sites, or slow replication in these mutants are unlikely causes of the preferential hyper distal USCE. Hence, we favor the model that Ctf4 and Ctf18 couple DNA replication and sister chromatid cohesion at the replication fork, thereby preventing the formation of aberant sister chroamatid structure that stimulates hyperdistal USCE.

Our USCE observations in yeast are likely to be relevant to the genomic stability in other eukaryotes. Like yeast, most if not all eukaryotes establish cohesion at many sites along the length of sister chromatids during S phase which are maintained until mitosis. The robustness of cohesion sites provides the potential to inhibit distal USCE as they do in yeast. Second, in our model USCE system, we measure USCE between a single repeat pair with only 589 bp of homology. However, most eukaryotic organisms have many more homologous repeats, such as transposable elements, along chromosome arms than budding yeast. Hence, the potential for distal USCE could be much greater, increasing the importance of its suppression by cohesin. Third, hemizygosity of genes encoding cohesin subunits or associated factors is strongly associated with cancer and developmental diseases (65). Cohesin hemizygosity could lead to increased cohesion site spacing and elevated USCE, which would generate duplications and deletions that are hallmarks of tumors. Fourth, the level of USCE caused by the hemizygosity could be further elevated by environmental conditions that cause replication stress mimicking *ctf4Δ* or *ctf18Δ* replication defects.

However, cohesin cannot be the sole repressor of USCE in yeast or other eukaryotes. First, in many species, proximal repeats are abundant in intergenic regions and introns. Proximal USCE between these repeats could lead to deleterious deletions within individual genes. Thus, repression of proximal as well as distal USCE would seem to be critical to maintain gene function. This inhibitory mechanism is likely to be cohesin independent since we failed to observe an elevation of proximal USCE in any of our mutants. Second, the cohesion site offset in mammalian cells can be as high as 100 kb rather than 10-20 kb in yeast (66–68), suggesting that mammalian cells may be more prone to distal repeat USCE. Thus, other mechanisms may also help suppress distal repeat USCE in mammalian cells. Given the overwhelming precedence for the relevance of yeast chromosome biology including cohesin to all eukaryotes, it will be informative to interrogate cohesin’s role in regulating USCE in other organisms.

## Acknowledgements

This work was funded by a National Institutes of Health grant, 1R35 GM-118189-06 (to D.K.) and NIH grant R35GM133678 (to W.M.) and a grant from the National Human Genome Research Institute (HG003143 to JD). J.D. is an investigator of the Howard Hughes Medical Institute.

## Methods

### Yeast strains, media and reagents

Yeast strains used in this study were A364A except for YLV11 and its derivatives which are W303 background (Table 3). YPD media was made as previously described (Guacci et al 1997). When required, Hygromycin B (Invitrogen cat# 10687010) was added to YPD media to a final concentration of 300μg/ml. Auxin (3-inoleacetic acid; Sigma-Aldrich cat#I3750) was made in DMSO as a 1M stock then added to media at a final concentration of 0.75μM to deplete AID tagged proteins. Pronase E (Sigma-Aldrich). Nocodazole (Sigma-Aldrich) was made as a 1.5mg/ml stock in DMSO then used at 15μg/ml in media to arrest cells in metaphase.

### Constructing strains for determining proximal and distal unequal sister chromatid exchange (USCE)

Table 3 provides the primers and plasmids used to generate wild-type and mutant strains used to measure USCE.

Wild-type strains to measure USCE on chromosome V. A 1.07kb fragment containing a 3’hyg fragment (last 2/3 of the Hygromycin orf + pTEF terminator) was generated by PCR of plasmid pAG32 using primers VG1124/VG1125. Wild-type haploid VG4187-1A was transformed with this 3’hyg fragment along with CRISPR plasmid pVG588 to integrate 3’hyg on chromosome V at 318kb, forming strain VG4214-10B.

**Table 2.**
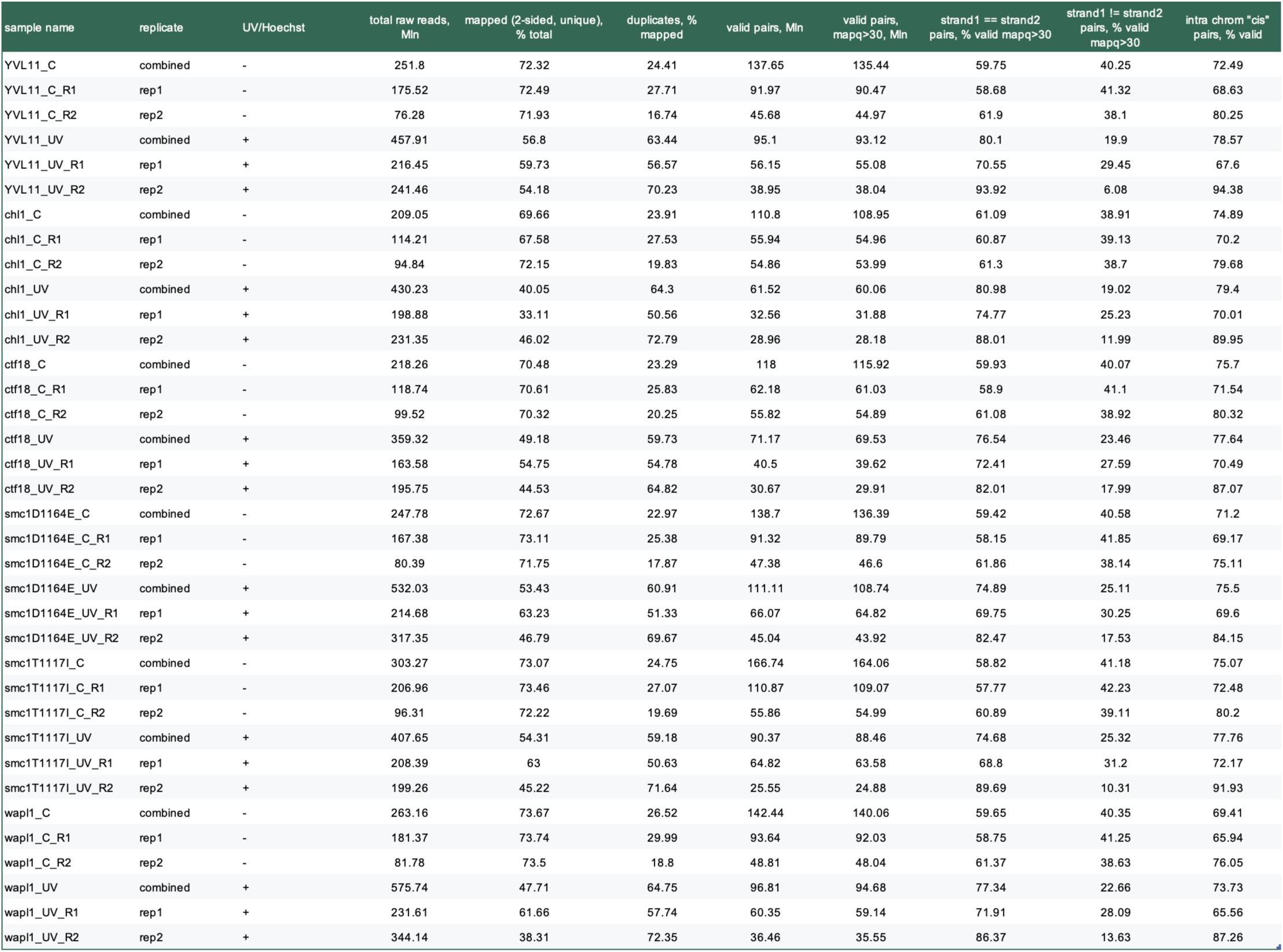

**Table 3.**

|  |  |
| --- | --- |
| VG4187-1A | *Mata leu2-3,112 ura3-52* |
| VG4214-10B | *Mata 3'hyg:ChrV-318kb leu2-3,112 ura3-52* |
| VG4220-14B | *Mata 3'hyg:ChrV-318kb 5'hyg:ChrV-322kb leu2-3,112 ura3-52* |
| VG4227-3B | Same as VG4220-14B except *wpl1Δ::NAT* |
| VG4231-7A | Same as VG4220-14B except *smc1-D1164E* |
| VG4235-5A | Same as VG4220-14B except *smc1-T1117I* |
| VG4239-9C | Same as VG4220-14B except *smc1-T1117W* |
| VG4249-7A | Same as VG4220-14B except *chl1Δ::G418* |
| VG4251-10A | Same as VG4220-14B except *ctf18Δ::G418* |
| VG4275-3B | Same as VG4231-7A except *smc3-K112R, K113R* |
| VG4280-8B | Same as VG4220-14B except *ctf4Δ0* |
| VG4259-13B | Same as VG4220-14B except *smc1-T1117I smc3-K113Q* |
| VG4423-4B | Same as VG4220-14B except *pol2-R567C* |
| VG4222-12D | *Mata 3'hyg:ChrV-318kb 5'hyg:ChrV-386kb leu2-3,112 ura3-52* |
| VG4228-4B | Same as VG4222-12D except *wpl1Δ::NAT* |
| VG4232-8C | Same as VG4222-12D except *smc1-D1164E* |
| VG4236-6B | Same as VG4222-12D except *smc1-T1117I* |
| VG4240-10B | Same as VG4222-12D except *smc1-T1117W* |
| VG4250-13D | Same as VG4222-12D except *chl1Δ::G418* |
| VG4252-18D | Same as VG4222-12D except *ctf18Δ::G418* |
| VG4260-15D | Same as VG4222-12D except *smc1-T1117I smc3-K113Q* |
| VG4258-5D | Same as VG4232-8C except *smc3-K112R, K113R* |
| VG4281-1D | Same as VG4222-12D except *ctf4Δ0* |
| VG4425-9D | Same as VG4222-12D except *pol2-R567C* |
| VG4330-3A | *Mata 3'hyg:ChrX-313kb leu2-3,112 ura3-52* |
| VG4460-13C | *Mata 3'hyg:ChrX-313kb 5'hyg:ChrX-317kb leu2-3,112 ura3-52* |
| VG4464-1A | Same as VG4460-13C except *ctf18Δ0* |
| VG4465-1D | Same as VG4460-13C except *smc1-D1164E* |
| VG4466-13B | Same as VG4460-13C except *wpl1Δ0* |
| VG4346-1A | *Mata 3'hyg:ChrX-313kb 5'hyg:ChrX-381kb leu2-3,112 ura3-52* |
| VG4362-2B | Same as VG4346-1A except *ctf18Δ0* |
| VG4363-4D | Same as VG4346-1A except *smc1-D1164E* |
| VG4364-5C | Same as VG4346-1A except *wpl1Δ0* |
| VG4275-11C | *Mata 3'hyg:ChrV-318kb 5'hyg:ChrV-322kb 5'hyg:ChrV-386kb leu2-3,112 ura3-52* |
| VG4279-9C | Same as VG4275-11C except *wpl1Δ::NAT* |
| VG4294-4B | Same as VG4275-11C except *ctf18Δ::G418* |
| VG4296-3A | Same as VG4275-11C except *smc1-D1164E* |
| VG3620-4C | *Mata LacO-NAT::lys4 GFP-LacI-HIS3:his3-11,15 TIR1-CgTRP1 leu2-3,112 ura3-52* |
| VG3902-3A | Same as VG3620-4C except *G418:MCD1-AID* |
| VG3956-14A | Same as VG3620-4C except *wpl1Δ::G418* |
| VG4138-2A | Same as VG3620-4C except *smc1-D1164E* |
| SX326 | Same as VG3620-4C except *chl1Δ0* |
| SX347 | Same as VG3620-4C except *ctf4Δ0* |
| VG4402-3C | Same as VG3620-4C except *ctf18Δ0* |
| VG4459-4C | Same as VG3620-4C except *pol2-R567C* |
| *YLV11 | *cdc21Δ::G418 trp1::TRP1-pGAL-dNK leu2::LEU2-pGAL-hENT1 a ura3-1 his3-11,15 can1-100* |
| *VG4308-4D | Same as YLV11 except *smc1-D1164E* |
| *VG4309-6A | Same as YLV11 except *ctf18Δ::G418* |
| *VG4312-1B | Same as YLV11 except *chl1Δ::NAT* |

**Table 4.**
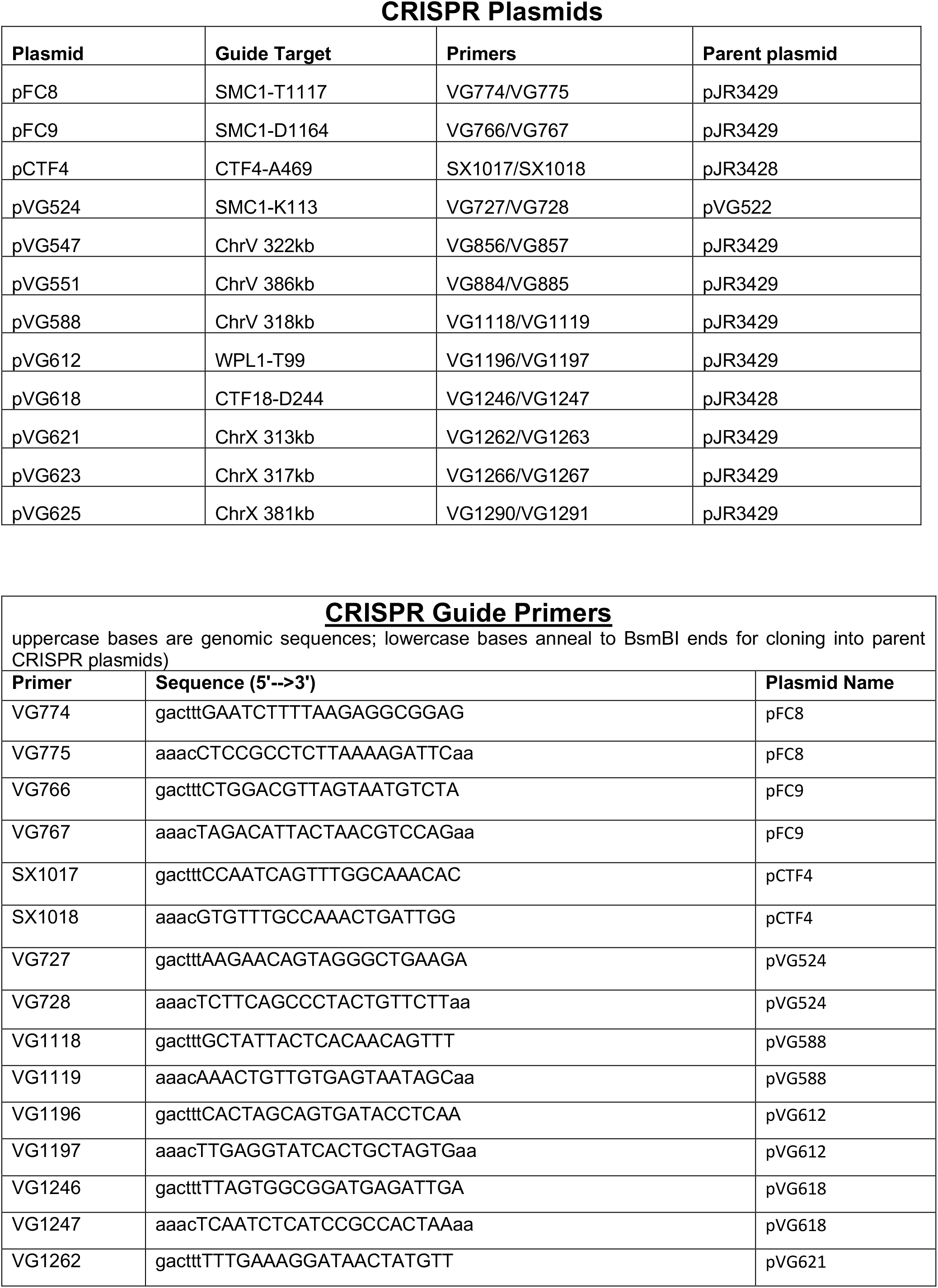
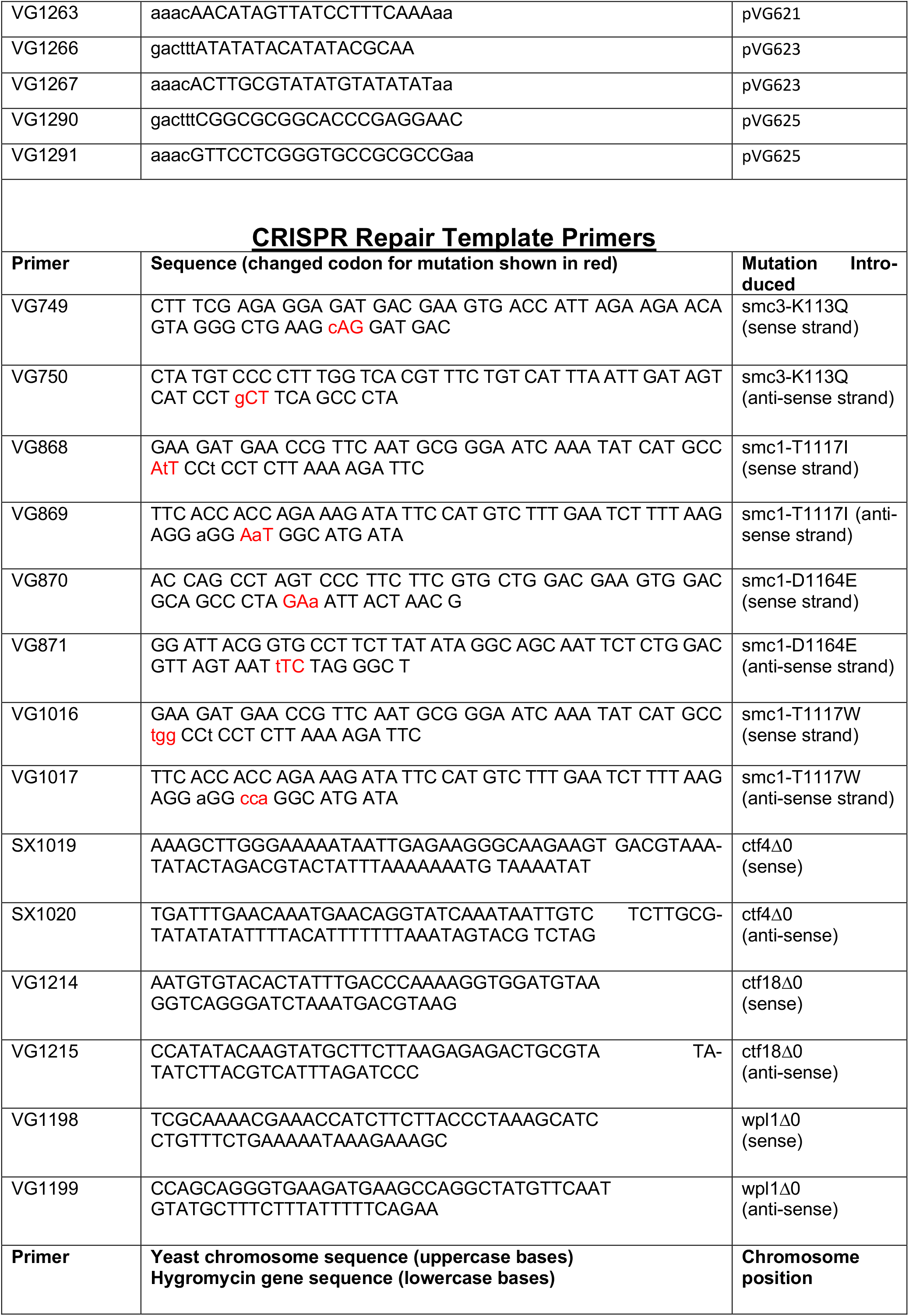
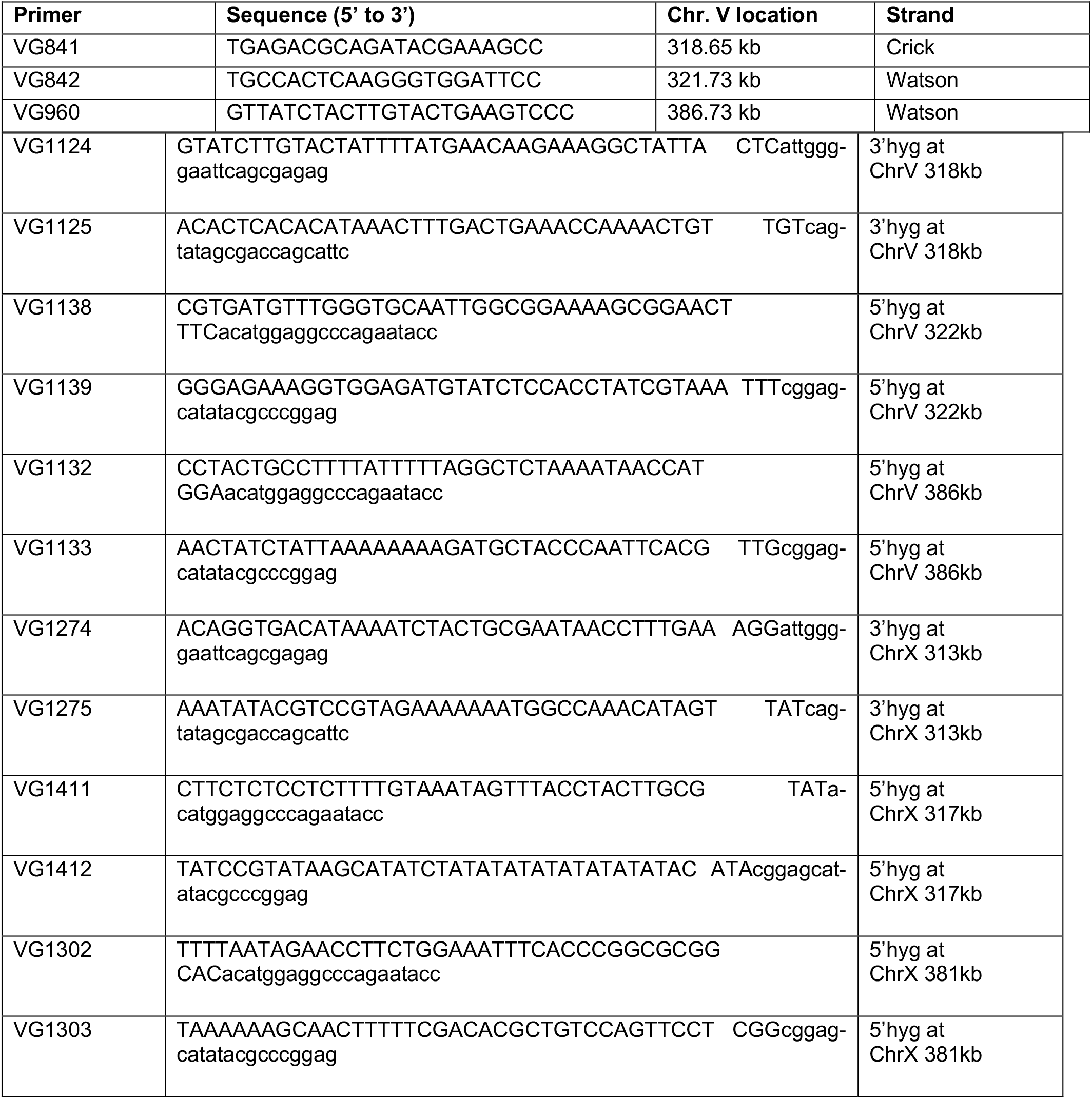

#### Proximal USCE

Wildtype strain VG4214-10B (w/ 3’hyg at 318kb) was transformed with a 1.23kb 5’hyg fragment (pTEF promoter + first 2/3 of the Hygromycin orf) generated by PCR of plasmid pAG32 using primers VG1138/VG1139 and CRISPR plasmid pVG547 to integrate 5’hyg on chromosome V at 322kb to form wild-type VG4220-14B.

#### Distal USCE

Wild-type VG4214-10B (w/ 3’hyg at 318kb) was transformed with a 1.23kb 5’hyg fragment (pTEF promoter + first 2/3 of the Hygromycin orf) generated by PCR of plasmid pAG32 using primers VG1132/VG1133 and & CRISPR plasmid pVG551 to integrate 5’hyg on chromosome V at 386kb to form wild-type VG4222-12D.

Wild-type strains to measure USCE on chromosome X. A 1.07kb fragment containing a 3’hyg fragment (last 2/3 of the Hygromycin orf + pTEF terminator) was generated by PCR of plasmid pAG32 using primers VG1274/VG1275. Wild-type haploid VG4187-1A was transformed with this 3’hyg fragment along with CRISPR plasmid pVG621 to integrate 3’hyg on chromosome X at 313kb, forming strain VG4330-3A.

#### Proximal USCE

Wildtype strain VG4330-3A (w/ 3’hyg at 313kb) was transformed with a 1.23kb 5’hyg fragment (pTEF promoter + first 2/3 of the Hygromycin orf) generated by PCR of plasmid pAG32 using primers VG1411/VG1412 and CRISPR plasmid pVG623 to integrate 5’hyg on chromosome V at 317kb to form wild-type VG4460-13C.

#### Distal USCE

Wildtype strain VG4330-3A (w/ 3’hyg at 313kb) was transformed with a 1.23kb 5’hyg fragment (pTEF promoter + first 2/3 of the Hygromycin orf) generated by PCR of plasmid pAG32 using primers VG1302/VG1303 and CRISPR plasmid pVG625 to integrate 5’hyg on chromosome V at 381kb to form wild-type VG4346-1A.

#### Mutants strains for analyzing USCE

Chromosome V USCE: mutants were inserted into wild-type strains VG4220-14B and VG4222-12D using CRISPR or standard molecular biology techniques.

Chromosome X USCE: mutants were inserted into wild-type strains VG4460-13C and VG4346-1A using CRISPR or standard molecular biology techniques.

All 3’hyg and 5’hyg fragment integrations and mutations inserted were confirmed by PCR sequencing and CRISPR plasmids lost before assessment of USCE.

### Determining the rate of unequal sister chromatid exchange (USCE)

Haploid strains bearing 3’hyg and 5’hyg fragments were inoculated into 5ml YPD then grown overnight at 30°C to saturation. Strains were dilution plated on YPD and grown 2d at 30°C to allow ∼50 well separated colonies /plate to form. Ten similar sized colonies from each strain were cut from YPD plates and added to separate eppendorf tubes containing 1 ml YPD. Tubes were vortexed to fully resuspend cells. Aliquots from each single colony were plated on hygromycin YPD media. At the same time a small aliquot of cells from each Eppendorf tube was pooled into a new eppendorf tube then dilution plated onto YPD. Plates were incubated at 30°C for 2 days. Cells which underwent uSCE grow into colonies on HYG plate. Colonies grown on YPD enable determination of average viable cells per colony. Rates of USCE were determined using the method of the median as described in Lea and Coulson (Genetics [1949] 49:264-285).

### Cohesion assay

Asynchronous cultures of cells were grown to mid-log phase at 30°C in YPD media, then alpha factor (alpha Factor) (Sigma) was added to 10^-8^M. Cells were incubated for 2h to induce arrest in G1 phase. Cells were synchronously released from G1 phase by washing 3 times containing 0.1 mg/ml Pronase E (Sigma) then resuspended in YPD containing nocodozale (Sigma) at 15ug/ml final and cells incubated at 30°C for 2h to arrest in mid-M (G2/M) phase. Cohesion was monitored at the *LYS4* arm locus as described (Boardman et al. 2023).

### Live cell imaging

Cell growth and preparation: Strains were grown to mid-log phase in YPD at 30°C. Alpha factor (Sigma-Aldrich) was added to cells at 10^-8^M final and cells incubated for 2h, Auxin (1mM final) was added after 1h. Cells were washed 3X in YPD containing Pronase E (0.1mg/ml final) then released YPD containing Pronase E (0.1mg/ml), Auxin (1mM final) and Nocodazole (15μg/ml final) and incubated for 80 minutes to arrest in G2/M. Cell cultures were concentrated 25-fold then resuspended in YPD + Auxin + Nocodazole. 200μl of cells were added to Concavalin A coated (see below) 14mm microwells of 35mm petri dish (Mat-Tek P35G-1.5-14-C) and incubated 15 minutes at room temperature to allow cells to settle and bind to the wells. Liquid was aspirated off then wells washed twice in imaging media (TRP-dropout media containing ethyl acetate (0.012% final), Nocodazole and auxin. 3.5ml imaging media was added to wells and imaging performed.

Concavalin A coated slides were prepared in advance as follows: Concavalin A (1mg/ml) was added to 14mm wells of the Mat-Tek 35mm petri dish and incubated 40 minutes on a Nutator to coat well bottom then aspirated off then washed once with water before use. Slide coating was done during nocodazole arrest. Imaging media was also prepared during nocodazole arrest and media placed on a nutator.

### SisterC

#### Cell Growth

Yeast Cell Cycle Synchronization for Sister-C Cell cultures were grown at 30°C in 50 ml YP media with 40 mg/L adenine, 100 µM thymidine and 2% galactose to midlog phase. Cells were then arrested in G1 with 5 μg/ml alpha factor peptide (Biomatik). 100 min after alpha factor addition 250 μM BrdU (Invitrogen B23151) was added. 130 min after alpha addition when 90% of cells were in G1 phase, cells were released from the block into YP media with 40 mg/L adenine, 500 μM BrdU and 2% galactose. Nocodazole (Sigma-Aldrich M1404) was added to cultures at 10 μg/ml 60 minutes after the release from G1. 4 hours after the G1 release cells were fixed with 3% formaldehyde (Fisher Scientific BP531) for 20 min at 30°C. 2.5 M glycine (Sigma-Aldrich 357002) was added for 5 min to quench the reaction. Cells were then washed and flash frozen in liquid nitrogen for sister-C processing.

#### SisterC library preparation

SisterC libraries were prepared as described previously (Oomen et al. 2020): Crosslinked cells were defrosted, washed and resuspended in 1 ml spheroplasting buffer (1 M Sorbitol, 50 mM Tris pH 7.5) with 5 µl of beta-Mercaptoethanol and 20 µg/ml 100T Zymolyase (USBio Z1004) for 10 min at 35°C. Cells were washed twice with 1x NEBuffer 3.1 (NEB B7203) and resuspended in 360 µl 1x NEBuffer 3.1. Chromatin was solubilized with 0.01% SDS for 5 min at 65°C followed by incubating on Ice and quenching with 1% Triton X-100. Chromatin was digested with 400U DpnII (NEB R0543) at 37°C overnight. After digestion, DpnII was inactivated by incubation for 20 min at 65°C. DNA ends were filled in using DNA polymerase I Klenow (NEB M0210) with nucleotides and biotin-14-dATP (Active Motif 14138) by incubating for 23°C for 4 hours in a ThermoMixer (900 rpm mixing; 10 secs every 5 mins). DNA fragments were ligated with T4 DNA ligase (Invitrogen 15224090) for 4 hours at 16°C in reactions of 75 μL each. All reactions were then combined and incubated overnight with proteinase K (Invitrogen 25530) at 65°C. DNA was purified using 1:1 phenol:chloroform followed by ethanol precipitation. Air-dried DNA pellets were resuspended in TLE (10 mM Tris-HCl, 0.1 mM EDTA, pH8) and incubated with 10 µg/ml RNAse A for 30 mins at 37°C before a 1X Ampure XP bead clean-up (Beckman Coulter A63881) was performed and DNA eluted in 100 µl TLE. Biotin was removed from unligated ends using T4 DNA polymerase (NEB M0203) with dATP and dGTP 20°C for 2 hours followed by enzyme inactivation at 75°C for 20 mins. DNA was sonicated to 600-800 bp using a Covaris M220 (Peak Incident power 50W, Duty cycle/factor 8%, 200 cycles, 24 seconds). Small DNA fragments were removed with a 0.6x Ampure XP bead capture, where the DNA captured by the beads was retained for the next steps. Sheard DNA ends were then repaired and A tailed using NEBNext Ultra II End Repair/dA-Tailing Module followed by NEBNext Adaptor ligation as per the manufacturers instructions (NEB E7546, E7595). 1X Ampure XP blead clean up was performed and DNA was eluted in 200 µl TLE. Each sample was split in two to obtain one Sister-C library treated with UV and Hoechst and one Hi-C library without treatment from the same biological sample. 100 ng/µl Hoechst 33342 (Invitrogen H3569) was added to each 100 µl Sister-C library and incubated for 15 min at RT in the dark. They were then UV irradiated at 2700 x 100 µJ/cm^2^. 1X Ampure bead clean-up was then performed on the UV samples. Biotin pull down was performed on both the Sister-C and Hi-C samples using MyOne Streptavidin C1 beads (Invitrogen 65001). Indexing was performed using NEBNext Multiplex Oligos for Illumina (NEB E7600) on the DNA attached to C1 beads; each Sister-C and Hi-C library was given a different i7 index. The final libraries were cleaned up using Ampure XP beads and eluted in 20 µl TLE. The libraries were sequenced using paired end 50 bp reads on an Illumina NextSeq 2000 system.

#### SisterC Analysis

Raw SisterC FASTQ files for all samples (non-UV controls and UV/Hoechst treated samples) were processed as previously described (Oomen et al. 2020) using the *distiller-nf* pipeline (v0.3.4) (Goloborodko et al. 2022). Briefly, reads were mapped to the sacCer3 reference genome using *bwa-mem* (Li 2013); subsequent parsing, deduplication, and filtering were performed with *pairtools* (Open2C et al. 2024), followed by data binning with *cooler* (Abdennur and Mirny 2020). Pairwise interactions (“pairs”) from UV-treated samples were classified following (Oomen et al. 2020): “along” (intra-sister) interactions were defined by alignments in the same orientation (“++” or “--”), while “between” (inter-sister) interactions were defined by opposite orientations (“+-” or “-+”). These classes were isolated using *pairtools select* with conditions “*strand1 == strand2*” and “*strand1 != strand2*”, respectively. Interaction data was binned at 1-kb and 2-kb resolutions and balanced via iterative correction (Imakaev et al. 2012), ignoring the first two diagonal bins. To ensure comparable biases between interaction classes, bias-correction weights derived from “along” samples were applied to their corresponding “between” counterparts, which was confirmed to yield flat corrected coverage. Individual replicate and combined sample statistics are provided in Table 2.

“Scaling plots,” which characterize the decay of interaction frequency (*P*) as a function of genomic distance (*s*), were generated from filtered pairs using the *scaling* module in *pairtools*. Sister interaction status (“along” vs. “between”) was assigned as described above.

Pileup plots anchored at Cohesin Associated Regions (CARs) and pairwise combinations of CARs were generated using the *pileup* function from *cooltools* (Open2c et al. 2024). This analysis utilized 1-kb binned interactions normalized by the distance-dependent decay of interaction frequency (“expected” values, calculated via the *expected_cis* function using sacCer3 chromosomal arms as the *view_df* parameter). The resulting plots were stratified by sister interaction status and genomic separation. CAR’s were adopted from (Costantino et al. 2020) as defined for the WT sample; notably, Wapl depletion demonstrated a redistribution of CAR strength without significant shifts in position.

In order to quantify the strength of such inter-CAR interactions, loop strength analysis calculations from (Akgol Oksuz et al. 2021) were followed. An enrichment of the center 3×3 area of the pileup was calculated relative to the average level of interactions in the periphery (average of the three 7×7 quadrants: upper left, upper right and lower right corners) see Supp Figure X. These enrichment values were calculated for each genomic distance range and plotted as a function of those genomic distances on Supp Fig X.

To characterize interactions between centromeres (CENs) and CARs, pileups were created using an “offset” normalization strategy as described in (Flyamer et al. 2017). For each CEN-CAR pair, the interaction was randomly offset along the CAR axis 40 times (20 upstream and 20 downstream shifts between 2 and 10 kb) while maintaining the CEN anchor. The resulting 40 snippets were averaged to normalize the input CEN-CAR snippet, thereby controlling for distance decay without interference from inter-chromosomal-arm interactions. CEN-anchored on-diagonal pileups were generated using 2-kb binned data after normalization by the chromosome-wide “expected.”

All code used for analysis is available as *Jupyter Notebooks* (Kluyver et al. 2016) on GitHub (https://github.com/dekkerlab/Koshland_collab.git). Abdennur, Nezar, and Leonid A. Mirny. 2020. “Cooler: Scalable Storage for Hi-C Data and Other Genomically Labeled Arrays.” Bioinformatics 36 (1): 311–316.

---

**Fig. S1.**
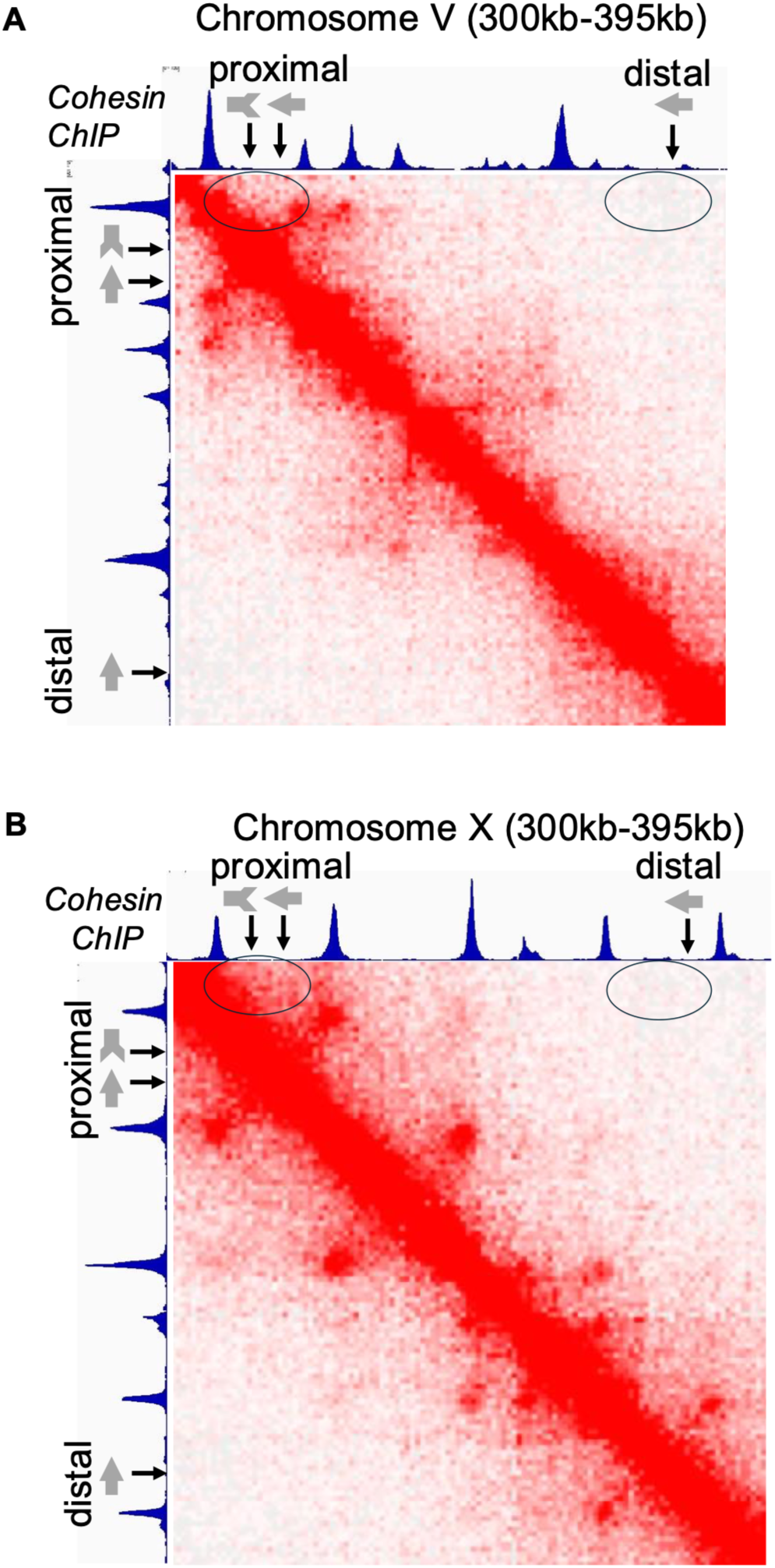
MicroC maps of chromosomes V and X showing where hyg fragments were inserted for uSCE assays and interactions in proximal regions or distal regions. (A-B) MicroC of WT yeast arrested in mid-M phase. A) Chromosome V right arm map from 300kb to 395kb with cohesin peaks defined by Mcd1p ChIP (blue peaks) Shown are the sites where the 3’hyg fragment (grey arrow tail) and the the 5’hyg fragments (grey arrowhead) at proximal (4kb) or distal (68kb) site, respectively. MicroC map of this region (red). Red dots off the main red axis show where adjacent DNA sites containing cohesin peaks interact thereby looping out intervening DNA. The black oval on the upper left shows interactions of DNA within the proximal region. The black oval on the upper right shows an absence of DNA interactions between the proximal and distal regions. B) Chromosome X left arm from 300kb to 395kb with hyg fragment insertion sites, Mcd1p ChIP and MicroC as described in (A). (C-D) MicroC of *wpl1Δ* yeast arrested in mid-M phase. C) Chromosome V as displayed as described in (A). D) Chromosome X as displayed as described in (B).

**Fig. S2.**
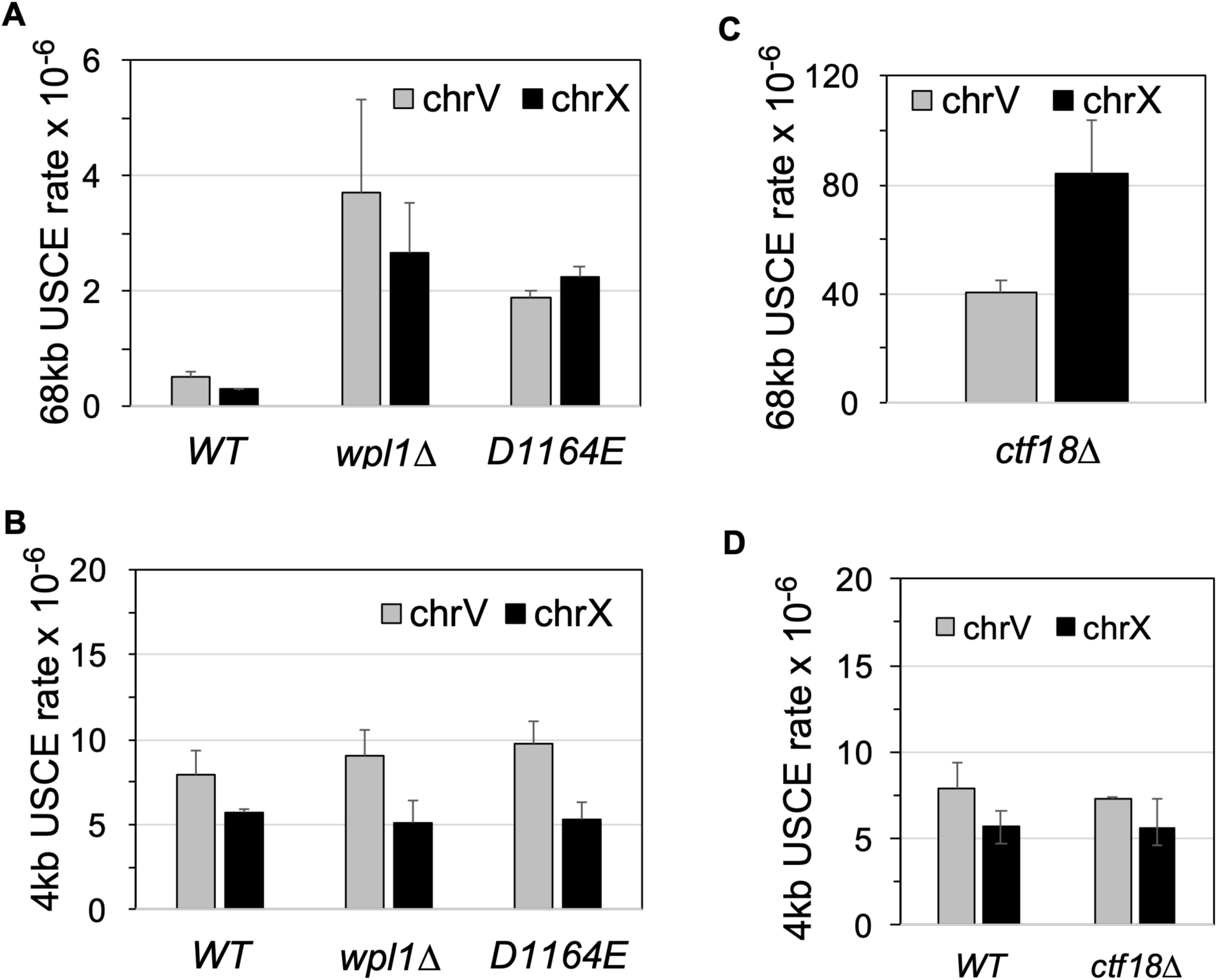
Mutants have similar USCE rates on chromosomes V & X. USCE rates for chromosome V (grey bars) and for chromosome X (black bars) are shown (A & C) Distal USCE for chromosomes V & X. Chromosome X uSCE data was generated using strains VG4346-1A (wild-type; WT), VG4364-5C (*wpl1Δ*), VG4363-4D (*smc1-D1164E*; DE) and VG4362-2B (*ctf18Δ*) as described (Materials and methods). Data for chromosome V was taken from Figures 3B, 3C & 6A and replotted for comparative purposes. (B) Proximal USCE for chromosomes V & X. Chromosome X uSCE data was generated using strains using strains VG4460-13C (wild-type; WT), VG4466-13B (*wpl1Δ*), VG4463-4D (*smc1-D1164E*; DE) and VG442-2B (*ctf18Δ*) as described (Materials and methods). Data for chromosome V was taken from Figures 3C and 6B and replotted for comparative purposes. Data for chromosome X was from 2 independent experiments.

**Fig. S3.**
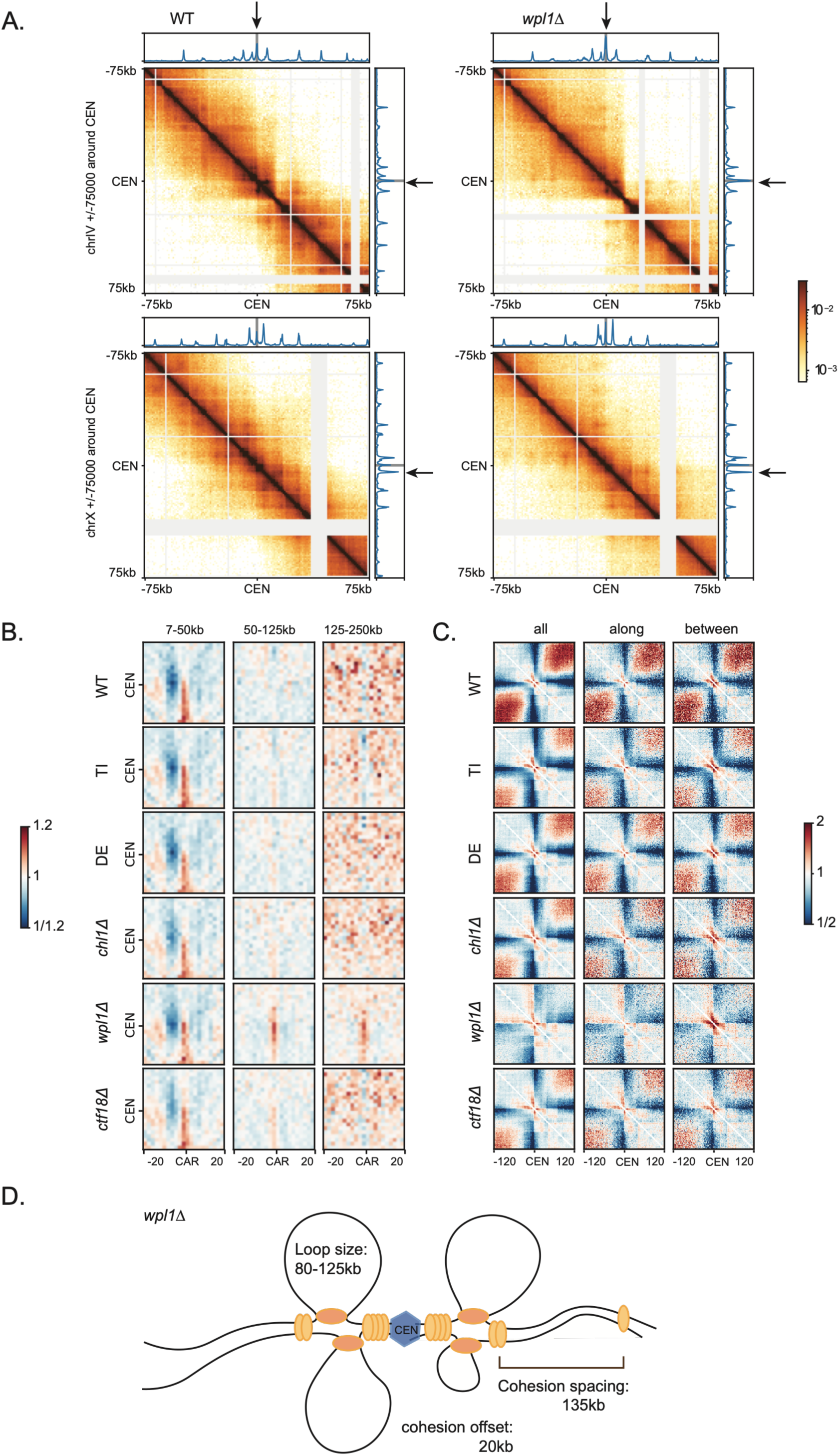
Large loops in *wpl1Δ* arise from centromeric cohesin. (A) Representative Hi-C maps plotted at 1kb resolution with Rad21 chip-seq tracks from chromosomes 4 and 10 in wild-type and *wpl1Δ* cells. The arrows indicate centromeric cohesin loops visible in *wpl1Δ* compared to WT. (B) Pile-up interaction maps based on one cohesin chip signal at the centromere with all arm CARs; plotted by distance from the centromere at a 2kb resolution. (C) peri-centromeric pile-ups of all interactions, intra-sister [along] interactions, and inter-sister [between] interactions. Plotted at a 2kb resolution. (D) Schematic representing the effects on sister chromatids of cohesive and extruding cohesin in a *wpl1Δ* background. Chip-seq of *wpl1Δ* showed increased binding of cohesin around centromeres. The centromeric and pericentromeric CARs appeared to have increased interactions with arm CAR’s showing visible spots on the heatmap (black arrows). When we piled up these interactions based on one centromeric CAR with all arm CAR sites we found that all the strains tested had interactions from the centromeric CAR to arm CAR sites up to 50 kb as expected. However, *wpl1Δ* had strong visible interactions up to 250 kb whereas no other strains we looked at had these interactions (Supp Figure NM B). We also noted an increase in trans cen-cen interactions in the *wpl1Δ* strain perhaps indicating strong cohesion at the centromere compared to the arm regions (supp Figure NM C). Overall, most cohesin being anchored at the centromeres, keeping centromeres cohesed, while being depleted along chromosome arms.

**Fig. S4.**
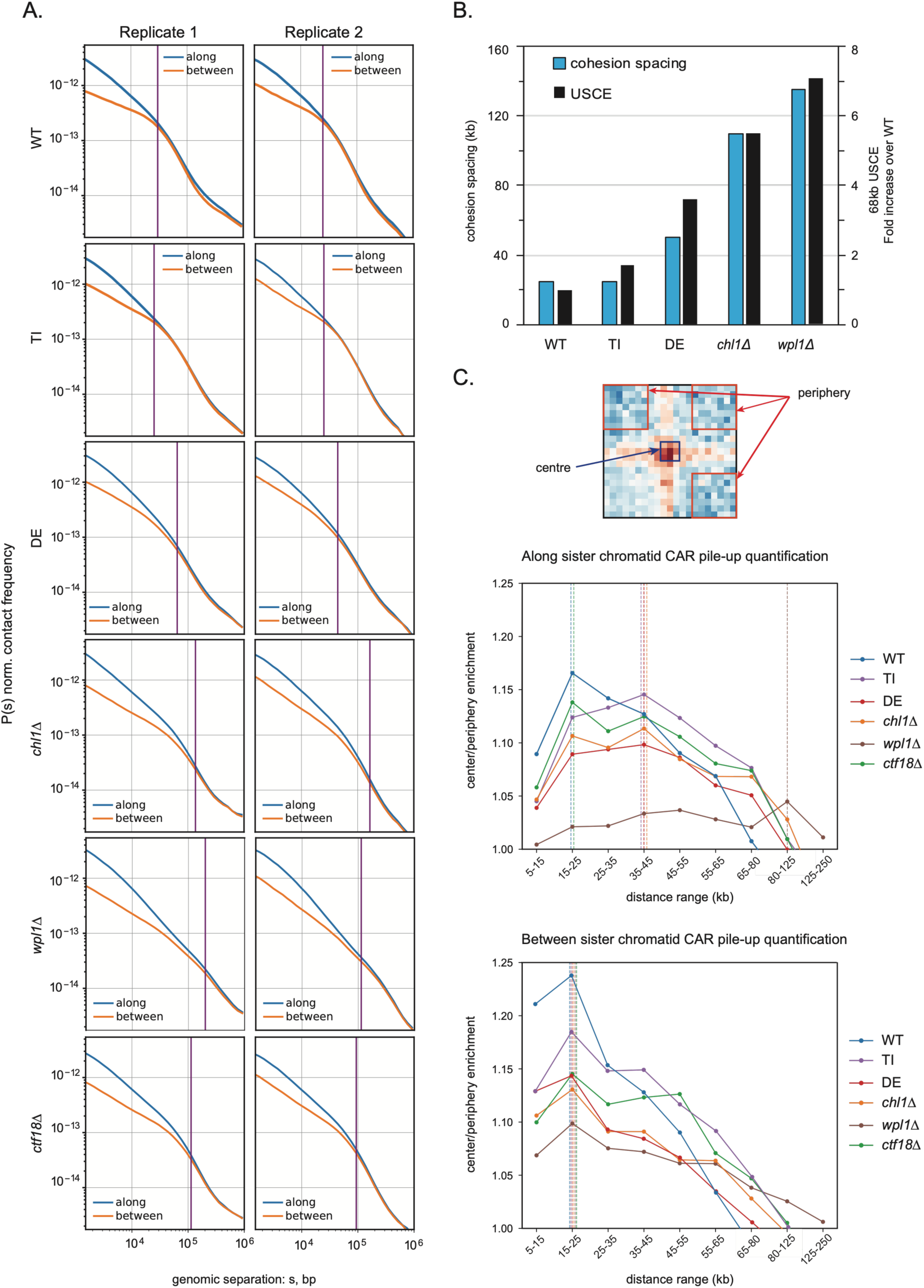
Sister-C replicates and quantification. (A) Contact frequency *P* plotted as a function of genomic separation for interactions either along or between sister chromatids. Two replicates shown that were combined in Figure 4A. (B) Dual axis histogram showing the 68 kb USCE fold increase (secondary y-axis) against cohesion spacing (primary y-axis) (from Figure 4A) for WT, TI, DE, *chl1Δ* and *wpl1Δ*. (C) Quantification of Figure 4B and 4C pile-up interaction maps. Inter-CAR interaction enrichment was calculated by normalising the average of the centre 3×3 pixels with the average of the peripheral 7×7 pixels from the three quadrants shown. Enrichment values for each distance shown for each strain. Dotted lines represent the strongest enrichment peak for each strain.

**Fig. S5.**
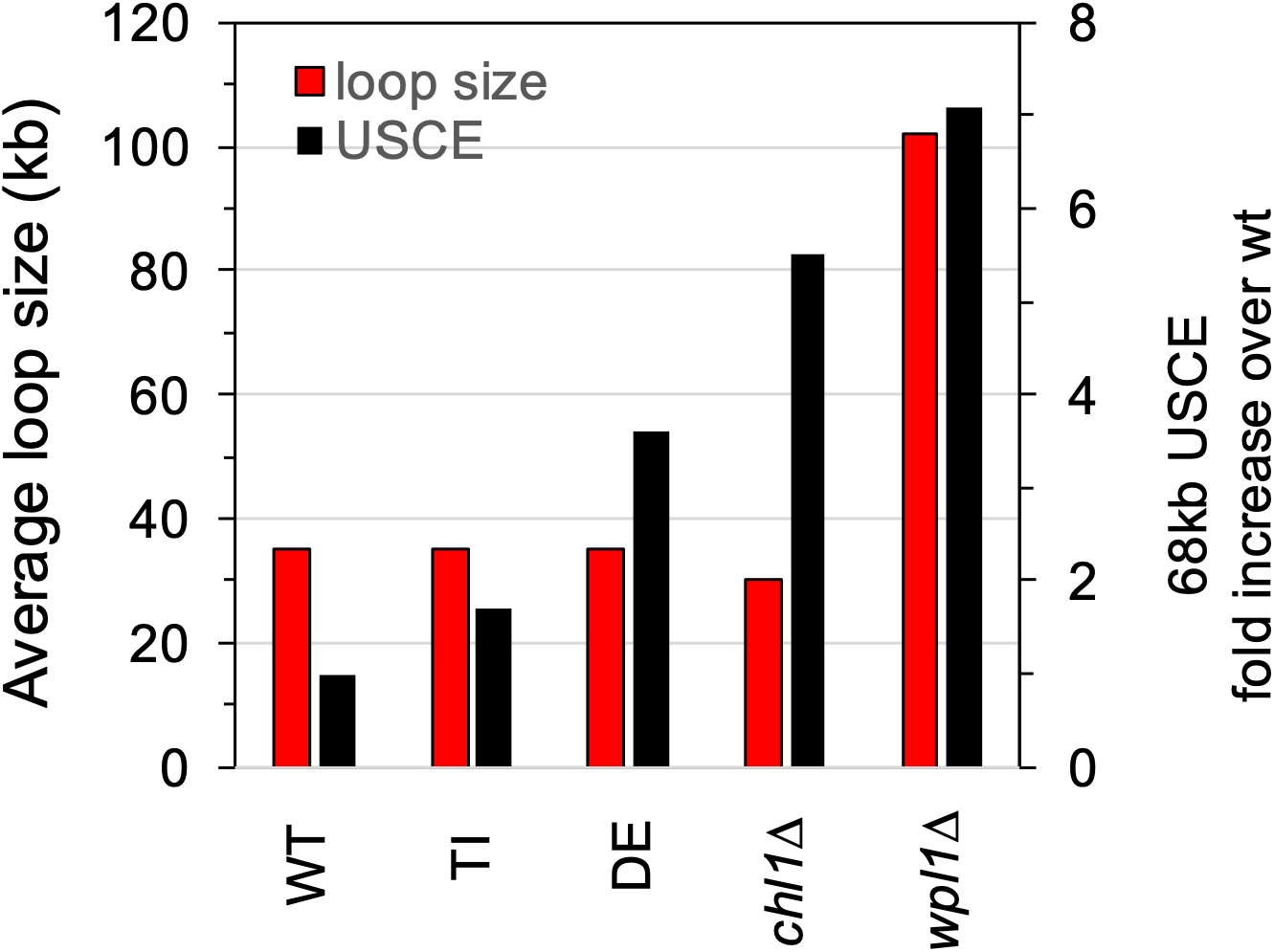
Loop size does not correlate with increased distal USCE rates seen in cohesin mutants. Comparison of average loop size of haploid wildtype strain [WT] and mutant haploid*s* [TI], [DE], [*chl1Δ*] and [*wpl1Δ*] taken from Fig. 4D plotted against the fold increase of distal USCE in each mutant taken from Fig. 3A calculated by dividing the rate of distal USCE in mutants by the rate for WT.

**Fig. S6.**
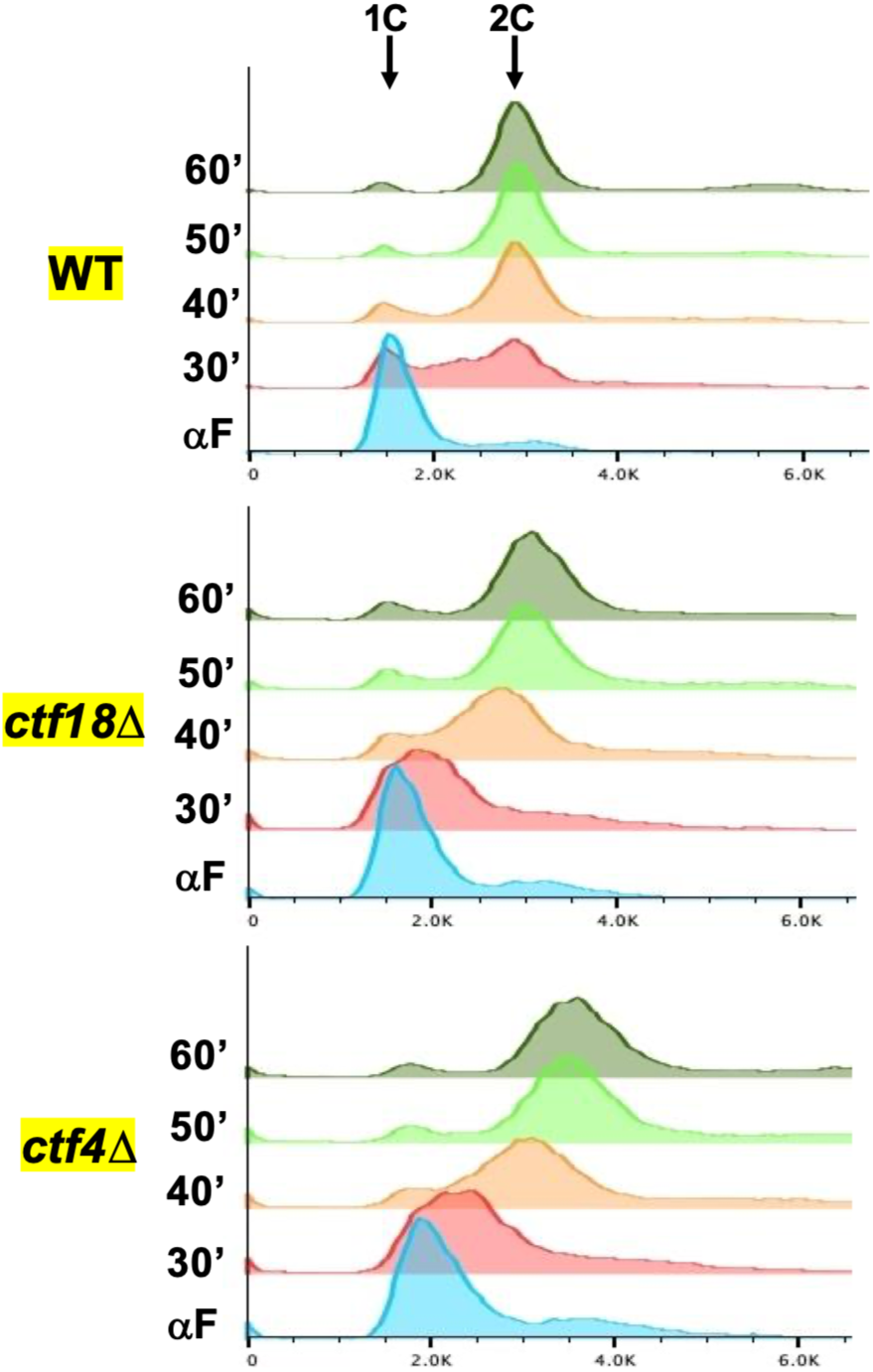
*ctf4Δ* and *ctf18Δ* mutants exhibit a DNA replication delay. Haploid WT (VG3620-4C), *ctf4Δ* (SX347) and *ctf18Δ* (VG4402-3C) were arrested in G1 phase and synchronously released from G1 and arrested in mid-M phase as described in materials and methods, except arrest was for 3h instead of 2h. Cells were fixed and processed for FACS (Materials and methods).

**Fig. S7.**
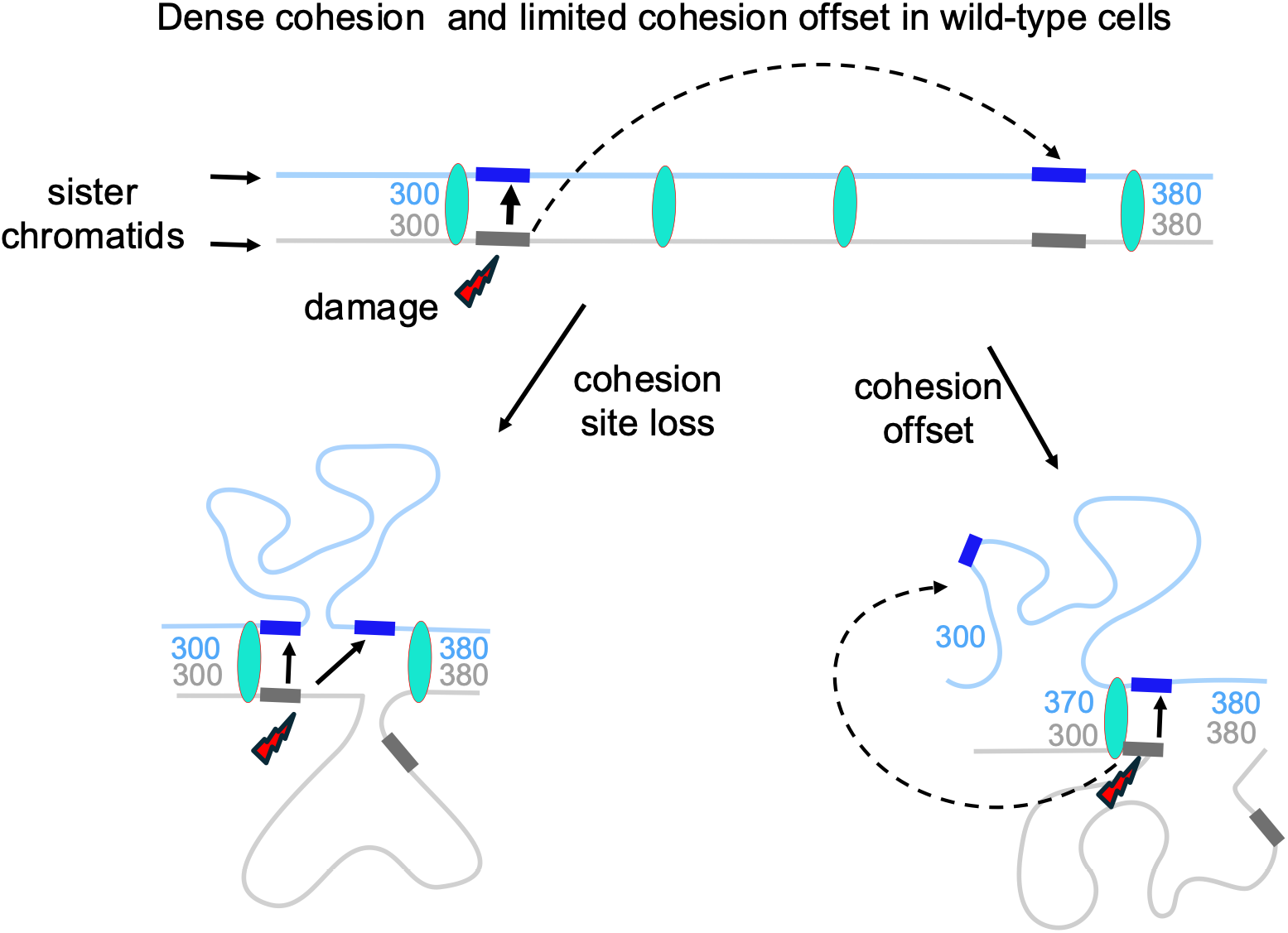
Model for cohesin dependent suppression of distal USCE. A pair of sister chromatids (blue and grey lines) are held together by cohesin (green oval). Blocks represent a homologous repeat sequence present at 300 and 370 kb. Damage to the grey repeat at 300 kb is repaired primarily using the blue repeat at the 300 kb (solid arrow) because this repeat is held in close proximity to the damaged repeat by cohesin (top). Dense cohesion also limits loop extrusion, preventing cohesin from bringing together the damaged repeat with the 370 kb repeat (not shown). Loss of cohesion sites allows the repeats at 300 kb to diffuse away from each other (bottom left). This improves chances for damaged 300 kb repeat to reach the repeat at 370 kb either diffusion or cohesin-dependent loop extrusion (hashed green oval). Alternatively cohesion offset occurs between sequences 300 kb on the grey sister chromatid and 370 kb of the blue sister chromatid (bottom right). When the repeat at 300 kb is damaged it is preferentially repaired by the repeat at 370 kb due to its greater proximity.

